# The terminal heme synthetic enzyme, Coproheme Decarboxylase, coordinates heme synthesis and uptake in response to iron in *Mycobacteria*

**DOI:** 10.1101/2021.05.10.443464

**Authors:** Rebecca K. Donegan, Jacqueline Copeland, Stanzin Edgha, Gabriel Brown, Owen F. Hale, Avishek Mitra, Hui Yang, Harry A. Dailey, Michael Niederweis, Paras Jain, Amit R. Reddi

## Abstract

Heme is both an essential cofactor and an abundant source of nutritional iron for the human pathogen Mycobacterium tuberculosis (Mtb). While heme is required for Mtb survival and virulence, it is also potentially cytotoxic. Since Mtb has the ability to both make and uptake heme, the de novo synthesis of heme and its acquisition from the host must be balanced in order to mitigate heme toxicity. However, the mechanisms employed by Mtb to regulate heme uptake, synthesis, and bioavailability are poorly understood. By integrating ratiometric heme sensors with mycobacterial genetics, cell biology, and biochemistry, we determined that the terminal heme biosynthetic enzyme, coproheme decarboxylase (ChdC), plays a role in regulating both heme bioavailability and uptake in Mtb. Moreover, we found that Mtb has a preference for scavenging reduced ferrous heme and exhibits a cell surface heme reductase activity that is regulated by ChdC. In Mtb, ChdC expression is down-regulated when iron is limiting, which in-turn increases both heme import and bioavailability. Such a mechanism may serve to protect cells from heme toxicity while trying to meet the nutritional demand for iron. Our results demonstrate that heme synthesis and uptake are tightly integrated in mycobacteria and represent the first example of a heme synthetic enzyme playing a role in controlling heme uptake.

**Significance Statement:** Heme is an essential but potentially cytotoxic cofactor and iron source for the pathogen, *Mycobacterium tuberculosis* (Mtb). To understand how Mtb coordinates heme uptake and synthesis to mitigate heme toxicity, we integrated heme sensors with mycobacterial genetics and biochemical approaches to probe the interplay between heme synthesis and scavenging. We discovered that the terminal heme synthetic enzyme, coproheme decarboxylase (ChdC), negatively regulates heme uptake and utilization in response to iron availability through a mechanism involving control of a ferric heme reductase. During iron limitation, ChdC is downregulated, thereby enhancing exogenous heme reduction, uptake and utilization while simultaneously suppressing heme synthesis, which allows Mtb to avoid heme toxicity. Our results highlight the close coordination between heme synthesis and uptake in mycobacteria.

**Classification:** Biological sciences : Biochemistry

## Introduction

Heme is both an essential cofactor and a primary source of iron for the human pathogen *Mycobacterium tuberculosis* (Mtb) (1, 2). As a cofactor, heme enables a number of physiological functions, including respiration, gas sensing, and protection against reactive oxygen and nitrogen species generated by the host immune system. Nutritionally, heme is the most bioavailable source of iron in the human host, with more than two-thirds of iron in circulation found bound to hemoglobin as heme-iron (3). Although heme is essential for Mtb, which can both make and scavenge heme, it is also potentially cytotoxic if present in excess or mishandled by cells (4, 5). As a consequence, Mtb must tightly regulate heme synthesis, import, and bioavailability in order to mitigate heme toxicity. However, the mechanisms underlying the regulation of heme homeostasis in mycobacteria are poorly understood (1, 6, 7).

At least two unique heme uptake pathways have been identified in Mtb; one that requires albumin for heme uptake and one that is albumin independent.(7-9) The presence of multiple pathways for heme scavenging implies that heme uptake is of great importance for Mtb virulence. Indeed, the knockout of the albumin independent pathway alone decreases survival in the macrophage.(8) While the factors that govern heme uptake in Mtb are largely unknown, heme and iron uptake in Mtb are regulated independently.(10)

In addition to uptake, Mtb encodes a complete heme biosynthetic pathway that begins with the synthesis of 5-aminolevulinic acid (ALA) from glutamate via the C_5_ pathway (11). ALA is then metabolized to coproporphyrinogen III in 4 enzymatic steps which are common among eukaryotes. However, unlike eukaryotes and gram-negative bacteria, which utilize the canonical protoporphyrin dependent branch (PPD) for the three terminal heme synthesis steps, Mtb synthesizes heme through the coproporphyrin dependent branch (CPD).(11-13) In the PPD branch, coproporphyrinogen III is oxidatively decarboxylated to protoporphyrinogen IX, followed by oxidation to protoporphyrin IX and insertion of iron by ferrochelatase to create heme.(11) In the CPD branch utilized by Mtb, coproporphyrinogen III is first oxidized to coproporphyrin III, followed by ferrochelatase-catalyzed iron insertion to make coproheme III and finally decarboxylation to make heme. The coproporphyrinogen oxidizing enzyme and ferrochelatase are homologous in the two pathways, despite binding different substrates.(11) However the terminal enzyme in Mtb, coproheme decarboxylase (ChdC), is unique to the CPD branch. (11) The divergence of heme synthesis strategies in Mtb and the necessity of heme for Mtb survival has led to the consideration of targeting heme synthesis for anti-Mtb therapies.(6, 13, 14) However, it is unclear what the relative contribution of exogenously scavenged and endogenously synthesized heme is towards filling the bioavailable heme pool and meeting the metabolic demands of Mtb. Moreover, given that in some organisms, such as the eukaryote *Saccharomyces cerevisiae*, heme import is upregulated when heme synthesis is ablated, (15) it is not known what impact perturbing heme synthesis may have on heme uptake pathways in mycobacteria.

Herein, by integrating genetically encoded ratiometric heme sensors with mycobacterial molecular genetics and biochemical assays, we probe the mechanisms underlying the coordination of heme uptake, synthesis, and bioavailability in Mtb and *Mycobacterium smegmatis* (Msm). Our results establish that both Mtb and Msm maintain a reservoir of exchangeable bioavailable heme and that *de novo* synthesized heme is more bioavailable than exogenously supplied heme. Moreover, we determine that heme uptake and utilization is negatively regulated by the terminal heme biosynthetic enzyme, ChdC. Msm and Mtb strains with a knockout of *chdC* significantly increase both the uptake and utilization of exogenous heme in both Msm and Mtb. Furthermore, Mtb ChdC expression is proportional to iron availability. When Mtb is iron limited, ChdC expression is downregulated, which in turn increases the uptake and utilization of exogenous heme. Our results further indicate that Mtb has a preference for taking up reduced ferrous heme and exhibits a cell surface heme reductase activity that is inhibited by ChdC. Altogether, ChdC coordinates the synthesis and uptake of heme in response to iron availability, providing a means to prevent heme toxicity while trying to meet the nutritional demand for iron using heme uptake pathways.

## Results

### Characterization of heme bioavailability in *M. smegmatis* and *M. tuberculosis*

Total cellular heme can be considered as the sum of exchange inert and labile heme (LH) (1, 4, 5, 16). The majority of intracellular heme is exchange inert, corresponding to heme that is tightly bound to hemoproteins. A smaller fraction of intracellular heme is maintained as a LH pool that is *bioavailable* for heme dependent processes. To measure LH within mycobacteria, we incorporated and validated previously described genetically encoded LH sensors (16-20) in Msm and Mtb (Supplemental Appendix and **Fig. S1 and S2**). <u>H</u>eme <u>s</u>ensor <u>1</u> (HS1) is a tri-domain construct consisting of the heme binding protein cytochrome *b*_562_ (Cyt *b*_562_), fused to enhanced green fluorescent protein (eGFP), and monomeric Katushka red fluorescent protein 2 (mKATE2). The fluorescence of eGFP is quenched upon heme binding to Cyt*b*_562_, whereas the mKATE2 fluorescence is unaffected.(16) Thus, the eGFP/mKATE2 fluorescence ratio is inversely proportional to cellular LH.(16)

In WT Msm, both the high affinity prototype sensor HS1, which exhibits dissociation constants of 3 nM for ferric heme (*K*_D_ ^Fe(III)^) and ∼1 pM for ferrous heme (*K*_D_^Fe(II)^) (16, 17), and a moderate affinity sensor variant HS1-M7A, which exhibits *K*_D_ ^Fe(III)^= 2 μM and *K*_D_ ^Fe(II)^ = 25 nM, (16) exhibit a dose-dependent increase in sensor eGFP/mKATE2 fluorescence ratio in response to succinylacetone (SA), an inhibitor of the heme biosynthetic enzyme porphobilinogen synthase (**Fig. S1 a-b**).(21) In contrast, HS1-M7A,H102A, which cannot bind heme and serves as a control for heme-independent changes to sensor fluorescence, did not exhibit SA-dependent changes in eGFP/mKATE2 fluorescence ratios (**Fig. S1c**). Together, these data indicated that both HS1 and HS1-M7A were competent for sensing intracellular LH in Msm. By adapting previously established sensor calibration procedures to Msm (**Fig. S1 d-f**, see Supplemental Appendix) we estimated that the heme occupancies of HS1 and HS1-M7A were > 85% and ∼50%, respectively, making HS1-M7A ideally suited for measuring LH in Msm. In the avirulent Mtb strain mc^2^6230 (WT Mtb), the heme sensors HS1 and HS1-M7A likewise exhibited heme-dependent fluorescence responses and the control, HS1-M7A, H102A, did not **(Fig. S2 a-c)**. The *in situ* calibration of the sensors in Mtb **(Fig. S2 d-f)** revealed that HS1 is saturated with heme, like in Msm, and HS1-M7A is < 20% bound to heme, unlike in Msm where HS1-M7A is ∼50% bound to heme. LH was only completely depleted from HS1 after repeated exposure to 500 μM SA, which was applied every 72 hours for 13 days (**Fig. S2 b**). Both exogenous hemin chloride (referred to hereafter as heme) (**Fig. S2 a**,**c**) and hemoglobin (Hb) (**Fig. S2c**) significantly increased HS1-M7A detected LH, which indicated that scavenged heme contributed to the LH pool. The heme biosynthetic precursor ALA had little effect on the LH pool of Mtb (**Fig. S2c**), which indicated that heme synthesis and/or partitioning of synthesized heme into the LH pool was tightly regulated. The observation that Msm has a greater concentration of bioavailable heme than Mtb, as assessed by HS1-M7A heme loading, is not due to generally higher levels of heme in Msm. In fact, Mtb had > 3-fold higher heme per cell **(Fig. S2g)** or nearly 7 times more heme per milligram of protein than Msm (**Fig. S2h**). Therefore, the reduced LH levels of Mtb result from differences in heme speciation and buffering between Mtb and Msm.

### Exogenous vs endogenous heme utilization in *M. smegmatis*

Since mycobacteria can both synthesize and uptake heme,(11, 22) we sought to determine if there was a difference in the bioavailability and utilization of exogenously supplied versus endogenously synthesized heme. Towards this end, we generated a heme auxotrophic strain of Msm with a deletion of the first heme synthesis enzyme, glutamyl tRNA reductase (GtrR) (11). Δ*gtrR* Msm exhibited a heme auxotrophy (**Fig. 1a**, black open squares with cyan line), and supplementation with 5 µg/mL ALA, which restored heme synthesis (**Fig. 1b**, gray bars), rescued growth to that of WT Msm (**Fig. 1a**, orange triangles and black circles, respectively). However, Δ*gtrR* Msm supplemented with 50 µM exogenous heme exhibited a substantial growth defect (**Fig. 1a**, red diamonds) compared to WT Msm (**Fig. 1a**, black circles) and Δ*gtrR* Msm supplemented with ALA (**Fig. 1a**, orange triangles). This growth defect was not due to heme toxicity as WT Msm supplemented with 50 µM heme (**Fig. 1a**, blue squares) had similar growth rates to untreated cells (**Fig. 1a**, black circles). These growth data indicated that Msm did not utilize exogenous heme as efficiently as endogenously synthesized heme.

**Fig. 1.**
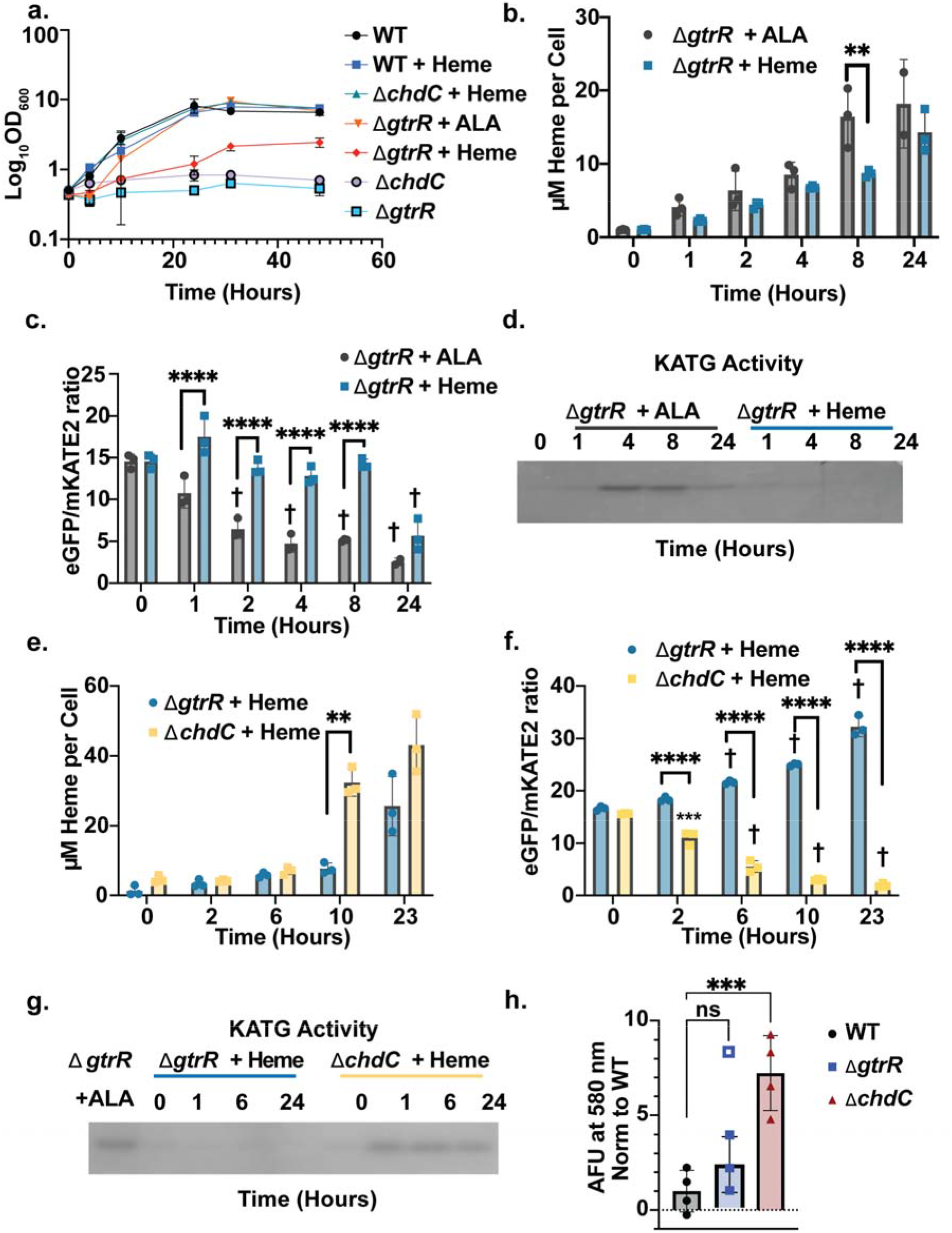
Utilization of exogenous versus endogenous heme in Msm is regulated by ChdC. (**a-d**) Effects of ALA (5 µg/mL) or heme (50 µM) supplementation on (**a**) growth rate, (**b**) total intracellular heme, (**c**) HS1-M7A detected labile heme, (**d**) and activity of the catalase-peroxidase KATG in Δ*gtrR* Msm cells. **e-g** Effects of heme (50 µM) supplementation on (**e**) total intracellular heme, (**f**) HS1-M7A detected labile heme, (**g**) and activity of the catalase-peroxidase KatG in Δ*gtrR* vs Δ*chdC* Msm cells. **h**. Uptake of the heme analog zinc mesoporphyrin (ZnMP) in WT, Δ*gtrR*, and Δ*chdC* Msm cells cultured with 1 µM ZnMP for 10 hours as quantified the ZnMP emission of ZnMP at 580 nm. Duplicates from two independent trials are shown. In panel **a**, growth curves represent the average cell density of triplicate cultures. In panels **b-c**, data represent the mean ± S.D. (error bars) of triplicate cultures. In panel **b**, the statistical significance was assessed by two-way ANOVA with Bonferroni post hoc test: ** *p* = 0.0015. In panel **c**, the statistical significance was assessed by two-way ANOVA with Bonferroni post hoc test. Black asterisks denote statistically significant differences at each time point: **** *p* <0.0001. Grey crosses denote statistically significant differences relative to “time 0” for each set of treatments. † *p* < 0.0001. In all panels, differences that are not statistically significant are unlabeled. The zymogram depicted in panel **d** is representative of 2 independent trials. In panels **e-f**, data represent the mean ± S.D. (error bars) of triplicate cultures. In panel **e**, the statistical significance was assessed by a two-way ANOVA with Bonferroni post hoc test: **** *p* < 0.0001, ** *p* = 0.0015. In panel **f**, the statistical significance was assessed by two-way ANOVA with Bonferroni post hoc test. Black asterisks denote statistically significant differences at each time point: **** *p* <0.0001. Grey crosses or asterisks denote statistically significant differences relative to “time 0” for each set of treatments: *** *p* = 0.0001; † *p* < 0.0001. In panel **h**, the statistical significance was assessed by one-way ANOVA with Dunnett’s post hoc test using WT Msm as control. For Δ*gtrR* the open square outlier was 1.4 standard deviations from the mean and was omitted from statistical analyses and ANOVA tests: ** *p* = 0.0009; “ns” denotes non-significant differences and *p* = 0.2178. In all panels, differences that are not statistically significant are unlabeled. The zymograms depicted in panels **d** and **g** are representative of 2 trials.

To determine if the growth disparity between endogenous and exogenous heme utilization corresponded to differences in intracellular heme accumulation and/or bioavailability, we measured total (**Fig. 1b**) and labile (**Fig. 1c**) heme in Δ*gtrR* Msm cells supplemented with heme or ALA. Δ*gtrR* Msm cells were starved of heme and ALA for 18 hours, which was sufficient to reduce total heme (**Fig. 1b**) and deplete LH (**Fig. 1c**). Δ*gtrR* Msm cells were then supplied with either 5 µg/mL ALA for endogenous heme synthesis, or 50 µM heme as a source of exogenous heme. Both ALA and heme supplementation restored total heme to a similar extent in Δ*gtrR* Msm cells over a broad time range (**Fig. 1b**). However, ALA more rapidly increased the bioavailable LH pool than exogenously supplied heme over the same time span (**Fig. 1c**). ALA supplementation resulted in a significant increase in LH within 2 hours, whereas it took more than 8 hours for exogenously supplied heme to have the same effect. Therefore, while both exogenous and endogenous heme contributed to intracellular heme accumulation, exogenous heme did not populate the LH pool as rapidly and was less bioavailable, thereby explaining its poor rescue of Δ*gtrR* cells.

The heme dependent catalase-peroxidase KatG plays an important role in detoxifying reactive oxygen species in mycobacteria and is important for mycobacterial virulence.(23) To determine if the difference in utilization of exogenous versus biosynthesized heme affected the activity of this hemoprotein, we measured KatG activity in Δ*gtrR* Msm using an in-gel catalase-peroxidase activity assay.(24, 25) We found that while endogenously synthesized heme rescued KatG activity, exogenous heme did not (**Fig. 1d**). Upon depletion of heme, Δ*gtrR* Msm did not have active KatG (**Fig. 1d**, time 0). Supplementation of Δ*gtrR* Msm with ALA yielded active KatG within 4 hours(**Fig. 1d**) However, heme supplementation did not yield active KatG even after 24 hours.(**Fig. 1d**) Altogether, these results revealed that while Δ*gtrR* Msm utilized exogenous heme for growth and metabolism, it was less bioavailable than endogenously synthesized heme.

### ChdC is a negative regulator of heme uptake and utilization in *M. smegmatis*

The terminal heme synthesis enzyme ChdC is unique to actinobacteria and firmicutes and had been put forth as a potential target for anti-Mtb therapies.(13) The poor ability of Δ*gtrR* Msm to utilize exogenously supplied heme led us to speculate that heme would be relatively ineffective at rescuing Msm lacking *chdC*. However, unlike in Δ*gtrR* cells, we found that Δ*chdC* Msm was fully rescued by exogenous heme (**Fig. 1a**, green triangles). To better understand the relationship between the heme synthetic enzymes GtrR and ChdC and heme uptake, bioavailability and utilization, we measured total heme, LH, and KatG activity in heme depleted Δ*gtrR* and Δ*chdC* Msm cells supplemented with exogenous heme. Heme supplementation resulted in a four-fold increase in total heme accumulation after 10 hours in Δ*chdC* cells compared to Δ*gtrR* cells, which suggested that ChdC may play a role in regulating heme uptake (**Fig. 1e**). Strikingly, unlike in Δ*gtrR* cells, exogenous heme resulted in substantially elevated LH in Δ*chdC* cells (**Fig. 1f**). At 2 and 6 hours after the addition of exogenous heme, when total heme is similar between Δ*chdC* and Δ*gtrR* Msm, LH was significantly elevated in Δ*chdC* Msm. Moreover, exogenous heme activated KatG in Δ*chdC* Msm, but not in Δ*gtrR* Msm (**Fig. 1g**). and Δ*chdC* cells accumulated ∼7-fold more of the fluorescent heme analog zinc mesoporphyrin (ZnMP) than WT and Δ*gtrR* cells, which internalized similar levels of ZnMP (**Fig 1h**, and **Fig S3a**). Taken together, our results indicated that heme biosynthetic enzymes play a substantial role in regulating the uptake and utilization of exogenous heme.

However, it was unclear if heme uptake and bioavailability was positively regulated by GtrR or negatively regulated by ChdC. To differentiate these possibilities, we assayed KatG activity, as a reporter of heme uptake and availability, in a Δ*gtrR* Δ*chdC* double knockout. The double knockout had KatG activity similar to Δ*chdC* cells when supplemented with exogenous heme (**Fig. S3b**,**c**), which implied that the increased heme utilization in Δ*chdC* Msm was due to the loss of ChdC, not the presence of GtrR. Moreover, these results also indicated that any potential build-up of heme biosynthetic precursors in Δ*chdC* cells did not contribute to the elevated heme uptake and utilization observed in Δ*chdC* cells. Additionally, the expression of catalytically inactive mutants of Mtb ChdC did not inhibit heme uptake (See Supplemental Appendix and **Fig. S4**), suggesting a role for ChdC-mediated coproheme decarboxylation, heme binding, or heme transfer to downstream targets in regulating heme uptake and utilization. Altogether, our results indicated that ChdC was a negative regulator of heme uptake in Msm.

### ChdC is a negative regulator of heme uptake and utilization in *M. tuberculosis*

To establish the role of ChdC as a negative regulator of heme uptake and utilization in Mtb, we generated heme auxotrophic Mtb strains with deletions of *gtrR* or *chdC* and tested their ability to utilize exogenous heme. Both Δ*gtrR* and Δ*chdC* Mtb exhibited a heme auxotrophy (**Fig. S5**). As with Msm, ALA rescued Δ*gtrR* Mtb cells more efficiently than exogenous heme. However, unlike Msm, Δ*gtrR* Mtb + ALA (**Fig. 2a**, green triangles) did not grow as efficiently as WT Mtb (**Fig. 2a**, purple circles), which suggested that Δ*gtrR* Mtb cannot import ALA in a manner that fully supports growth in the indicated culture conditions. Moreover, Δ*chdC* Mtb supplemented with exogenous heme (**Fig. 2a**, red diamonds) grew at a rate similar to WT Mtb and faster than heme supplemented Δ*gtrR* cells (**Fig. 2a**, orange triangles). Additionally, Δ*chdC* cells accumulated more ZnMP than WT and ΔgtrR Mtb cells (**Fig. 2b, c**), which internalized similar levels of ZnMP. The 48-hour timepoint where we observed a ∼50% increase in ZnMP uptake in Δ*chdC* cells compared to WT, was approximately two doubling times for Mtb. This timepoint corresponded to the 10-hour timepoint for Msm (**Fig. 1h**), which had a doubling time of approximately 5 hours. Similar to heme uptake in Δ*chdC* Msm (**Fig. 1b)**, early time points for Mtb ZnMP uptake did not exhibit increased ZnMP uptake in Δ*chdC* Mtb (**Fig. S6**).

**Fig. 2.**
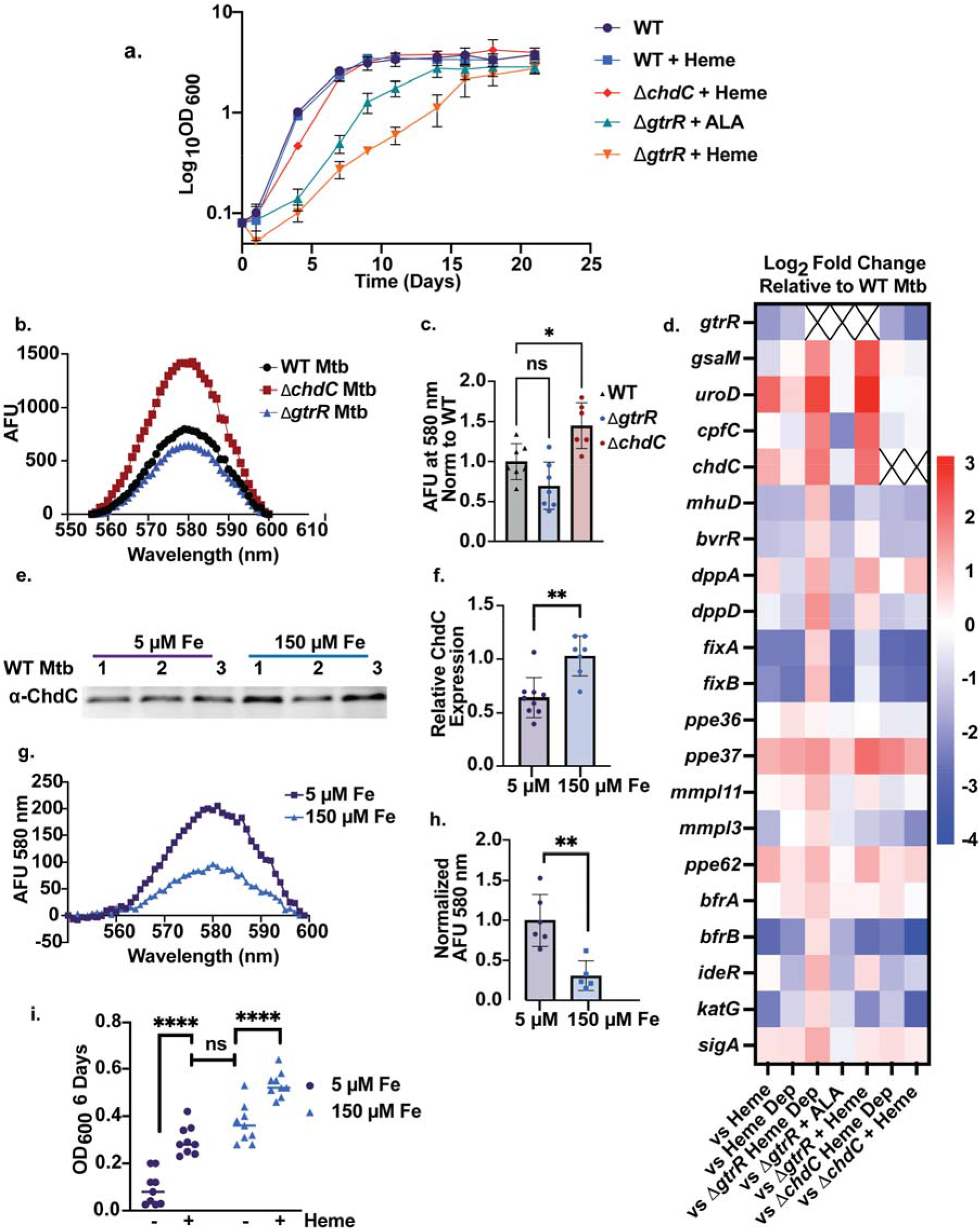
ChdC is a negative regulator of exogenous heme utilization in Mtb. (**a**) Effects of ALA (5 µg/mL) or heme (50 µM) supplementation on growth of WT, Δ*gtrR*, or Δ*chdC* Mtb strains. (**b**-**c**) Uptake of the heme analog zinc mesoporphyrin (ZnMP) in WT, Δ*gtrR*, and Δ*chdC* Mtb cells cultured with 1 µM ZnMP for 48 hours as measured by fluorescence. (**b**) Representative average of 3 ZnMP fluorescence spectra from one experimental trial and (**c**) quantification of the ZnMP emission peak at 580 nm from replicates in three independent trials are shown. **d**. Heat map of Log_2_ fold change of Mtb cultures compared to WT Mtb cultured in 7H9+ADS. Data shown is an average of two biological replicates. Xs represent Log_2_ fold change of < -6 for genetic knockouts. (**e**) Representative western blot of ChdC expression in WT Mtb cultured in low (5 µM Fe added) or high (150 µM Fe added) iron minimal media for 6 days. (**f**) Quantification of ChdC expression normalized to Ponceau S total protein stain from replicates in 3 independent trials n=9 for 5 µM and n=7 for 150 µM. (**g-h**) Fluorescence detected uptake of the heme analog ZnMP in WT Mtb cells cultured in low (5 µM) or high (150 µM) iron minimal media for 6 days after exposure to 1 µM ZnMP for 48 hours. (**g**) Representative ZnMP fluorescence spectra and (**h**) quantification of the ZnMP emission peak at 580 nm from replicates in two independent trials are shown, n=6 for 5 µM and n=5 for 150 µM. i. Effects of heme (12.5 µM) on growth of WT Mtb in minimal media (MM) supplemented with 5 µM or 150 µM Fe, n=9. In panel **a**, growth curves represent the average cell density of triplicate cultures. In panel **c** the statistical significance was assessed by a one-way ANOVA with Dunnett’s post hoc test using WT Mtb as control. In panel **c**, * *p* = 0.0152 and “ns” denotes a non-significant *p* = 0.0911. In panels **f** and **h**, the statistical significance was assessed by a two-tailed unpaired t-test: (**f**) ** *p* = 0.0010 and (**h**) *p* = 0.0023. In panel **i**, the statistical significance was assessed by using a two-way ANOVA with Bonferroni post hoc test: “ns” denotes a non-significant *p* = 0.2338; **** *p* < 0.0001; all unlabeled pairwise comparisons have *p* < 0.0001 but are omitted for clarity.

### Heme dependent transcriptional responses in *M. tuberculosis*

In order to assess the status of heme metabolism and utilization between *de novo* synthesized and exogenously scavenged heme, as well as ChdC-regulated heme uptake, we examined the expression of a wide-panel of heme homeostatic genes in response to heme in WT, Δ*gtrR*, and Δ*chdC* cells using RT-qPCR. Markers for heme metabolism included transcripts encoding genes for heme biosynthetic enzymes, *e*.*g. gtrR, gsaM, cpfC, urod* and *chdC*,(11) heme catabolizing enzymes *e*.*g. mhuD and bvr*,(26, 27) heme transport proteins, *e*.*g. ppe36, ppe37, ppe62, dppA, dppD* and *mmpl3*,(7-9, 28) additional factors previously implicated in heme uptake, *e*.*g. fixA, fixB, mmpl11*,(9, 10) iron storage proteins, *bfrA* and *bfrB*,(29) a heme-dependent peroxidase, *katG*,(30) and an iron homeostatic factor, *ideR*.(31) For heme treatments, WT, Δ*gtrR* and Δ*chdC* Mtb cells were cultured in 7H9 + albumin dextrose and salt (7H9+ADS, see methods) supplemented with 25 µM heme until growth reached saturation. Cells were then either washed and diluted into 7H9+ADS without heme for 3 days (“heme depleted”) or diluted into 25 µM heme for 3 days. Δ*gtrR* Mtb + ALA cells were cultured in 7H9+ADS supplemented with 5 µg/mL ALA. As a control, untreated WT Mtb cells were continuously cultured in 7H9+ADS and served as a reference to evaluate heme status in all mutants and growth conditions.

In WT Mtb, exogenous heme supplementation regulates the expression of the early and late enzymes of the heme biosynthetic pathway distinctly, with heme causing a 4-fold decrease in *gtrR* transcript and a 2.5-fold increase in *chdC* transcript (**Fig. 2d and Dataset S1)**. The differential effects of exogenous heme on *gtrR* and *chdC* expression suggests ChdC plays additional roles that extend beyond heme synthesis, including regulating heme uptake and bioavailability. Additionally, heme treatment of WT Mtb upregulated only the cell surface heme transport proteins *ppe37* and *ppe62*, while other heme uptake proteins (*ppe36, dppA, and dppD*) had little to no change in transcript, or in the case of *mmpl3*, had decreased expression. Exogenous heme downregulated the expression of the heme degrading heme oxygenase, *mhuD*, and the ferritin, *bfrB*, possibly indicating that heme degradation to release iron may not be occurring to an appreciable extent. Of note, WT Mtb cells depleted of exogenous heme showed lingering effects of heme exposure in their transcriptional profiles (**Fig. 2d**, column 2). This may be due to the continued internalization of recalcitrant heme retained at the cell surface even after washing and removal of exogenous heme from the cells and culture media.

Heme depleted Δ*gtrR* cells exhibited a transcriptional profile consistent with a heme starvation state, with all transcripts encoding heme biosynthetic and uptake proteins being induced (**Fig. 2d**). Supplementing Δ*gtrR* cells with ALA, but not exogenous heme, generated a heme replete state, with expression of heme synthesis and transport genes being repressed or restored relative to WT (**Fig. 2d**). The heme vs. ALA dependent transcriptional responses in Δ*gtrR* cells are consistent with endogenous heme being more bioavailable than exogenously supplied heme (**Fig. 2d and Fig. S7a**). Interestingly, restoring heme synthesis in Δ*gtrR* cells with ALA resulted in iron limitation, as inferred from the ∼3-fold decrease in *ideR* and *bfrB*, iron homeostatic genes that positively correlate with iron levels.(29, 32) Consistent with the notion that heme synthesis can limit iron, heme depleted Δ*gtrR* cells, which cannot synthesize heme, have a 2-fold increase in *ideR* and 1.5-fold increase in *bfrB*, indicating a more iron replete state relative to WT cells. Although exogenous heme does not rescue the heme deficiency of Δ*gtrR* cells, it does serve to repress *bfrB*, indicating heme can alter cellular iron homeostasis (**Fig. 2d and Fig. S7b**), albeit through an unknown mechanism. Surprisingly, *mhuD* expression did not correlate with heme levels as one might expect, with heme depleted Δ*gtrR* cells exhibiting elevated *mhuD* transcript and ALA or heme supplemented Δ*gtrR* cells exhibiting depressed *mhuD* levels. Altogether, transcriptomic profiling indicates that ALA, but not exogenous heme, effectively alleviates the heme deficiency of Δ*gtrR* cells, and that heme synthesis, and exogenous heme to a certain degree, can contribute to limiting cellular iron.

Surprisingly, heme depleted Δ*chdC* cells do not exhibit many of the transcriptional hallmarks of heme starvation observed in heme depleted Δ*gtrR* cells, including the induction of heme biosynthetic and transport genes (**Fig. 2d**). In fact, heme depleted Δ*chdC* Mtb had a similar transcriptional response to heme depleted WT Mtb, with one exception being *mmpl*3, which had a > 2-fold reduction in Δ*chdC* cells compared to WT (**Fig. 2d**). Transcript levels in heme treated Δ*chdC* cells were very similar to heme treated WT cells (**Fig. 2d and Fig. S7c**). One exception was the regulation of the heme biosynthesis gene *uroD*, which was upregulated in WT Mtb treated with heme but did not have altered transcription in heme treated Δ*chdC* Mtb (**Fig. 2d**). Given that *chdC* and *uroD* are predicted to be in the same operon, it is possible the loss of *chdC* alters expression of *uroD* through an undiscovered regulatory mechanism or due to differences in intracellular heme in Δ*chdC* cells compared to WT cells. Additionally, the transcript levels of the iron homeostasis genes *ideR* and *bfrB* were decreased ≥ 2-fold in heme treated Δ*chdC* compared to heme treated WT Mtb (**Fig. 2d** and **Fig. S7c**). Since *bfrB* is repressed in response to exogenous heme in WT Mtb (**Fig. 2d**), the more severe reduction of *bfrB* in heme treated Δ*chdC* cells compared to WT is consistent with the observation that heme uptake is increased in Δ*chdC* cells. Notably, none of the previously characterized heme uptake proteins were strongly induced in heme depleted Δ*chdC* cells relative to WT or Δ*gtrR* strains, indicating that ChdC regulation of heme uptake and bioavailability does not occur through the transcriptional regulation of known heme transport proteins. Altogether, the qPCR data further highlights that in Mtb exogenous heme is metabolized differently than endogenously synthesized heme.

### Iron availability regulates ChdC expression and heme uptake in *M. tuberculosis*

Since ChdC was found to be a negative regulator of heme uptake and utilization, we sought to model conditions experienced by Mtb during infection that may increase or decrease ChdC expression and in turn alter heme uptake and/or utilization. Given that heme can be used as an iron source under iron depleted conditions,(22) iron availability is necessary for heme synthesis, and ablating *chdC* increased heme uptake, we examined whether iron availability affected ChdC protein levels and heme uptake. Through immunoblotting experiments using a polyclonal anti-Mtb ChdC antibody, we determined that ChdC expression was decreased under low iron conditions in minimal media (MM) supplemented with 5 µM Fe (**Fig. 2 e,f**) compared to MM with 150 µM Fe.

To determine whether iron availability affected heme uptake in Mtb, we measured uptake of the heme analog ZnMP in Fe limited (5 µM Fe) and Fe replete (150 µM Fe) conditions. WT Mtb cells grown in MM supplemented with 150 µM Fe had ZnMP uptake that was ∼30% of WT Mtb grown in 5 µM Fe (**Fig. 2 g,h**).

Finally, we assessed the ability of WT Mtb to use exogenous heme for growth in iron limiting conditions that correspond to decreased ChdC expression (MM + 5 µM Fe). The addition of 12.5 µM heme increased WT Mtb growth in MM supplemented with 5 µM Fe and to the same level as cells treated with 150 µM Fe (**Fig. 2i**). Therefore, iron dependent ChdC expression negatively correlated with both an increase in the uptake of heme analog ZnMP and the effect of heme supplementation on WT Mtb growth in MM.

### Exogenous heme decreases porphyrin synthesis in *M. tuberculosis*

Heme synthesis is regulated by intracellular heme levels in some bacteria (11) and the decrease of *gtrR* transcript levels in WT Mtb with heme treatment (**Fig. 2d**) suggested that this regulation may occur in Mtb as well. To better determine if exogenous heme provides feedback to regulate heme synthesis in mycobacteria, we measured fluorescent porphyrins as a proxy for flux through the heme biosynthetic pathway. The results showed that exogenous heme decreased porphyrin levels in Mtb similar to inhibition of heme synthesis by succinylacetone, suggesting that heme does regulate heme synthesis early in the biosynthetic pathway. (Supplemental Appendix and **Fig. S8a-h**) Additionally, culturing WT Mtb in iron deplete MM (5 µM Fe) did not reduce porphyrin or heme levels compared to iron replete MM (150 µM Fe) **(Fig. S8 i**,**j)**. Taken together with the RT-qPCR results (**Fig. 2d**), we propose that heme feeds back to regulate the synthesis of early porphyrin intermediates and that this feedback likely requires *gtrR*.

### ChdC regulates a reductive heme uptake pathway in *M. tuberculosis*

In order to determine biochemical features of ChdC-regulated heme uptake, and ultimately its mechanism of action, we assessed if ChdC inhibited heme uptake in an oxidation-state dependent manner. Towards this end, we first assessed if Mtb exhibited a preference to internalize divalent zinc(II)protoporphyrin IX (ZnPP) or trivalent gallium(III)protoporphyrin IX (GaPP), which are redox inactive fluorescent heme analogs that serve as mimics of ferrous (Fe^2+^) or ferric (Fe^3+^) heme, respectively, due to similar metal oxidation numbers and ionic radii. (33-37) In standard growth conditions of 7H9 + ADS, ZnPP accumulated in WT Mtb to a far greater extent (nearly 3-fold in 48 hours) than GaPP (**Fig. 3a**). We next compared GaPP vs. ZnPP uptake in WT, Δ*gtrR* and Δ*chdC* Mtb in 7H9+ADS after 48 hours. In Δ*chdC* Mtb, the ratio of GaPP to ZnPP uptake increased ∼ 3-fold and approached 1 (**Fig. 3b**), which indicated that Δ*chdC* cells lost the ability to discriminate between the Fe^3+^-heme mimic GaPP and Fe^2+^-heme mimic ZnPP.

**Figure 3.**
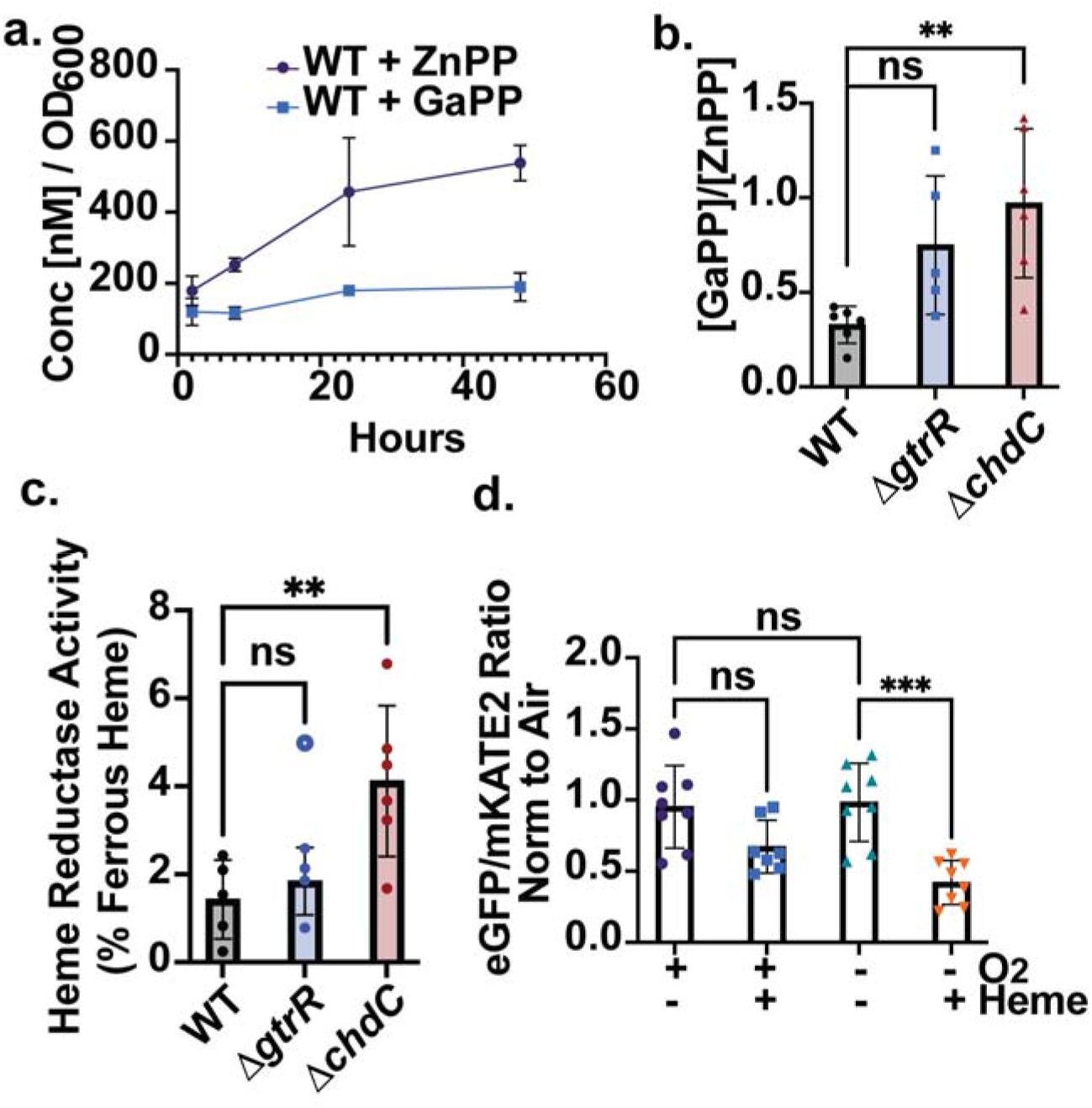
Mtb ChdC regulates reductive heme uptake. **a**. Binding and uptake of ZnPP and GaPP by WT Mtb measured over time by whole cell fluorescence. Concentrations were determined by standard curve. Data shown is from biological triplicates. **b**. GaPP/ZnPP ratio of WT, Δ*gtrR* and Δ*chdC* Mtb at 48 hours from two independent experiments. **c**. Percent ferrous heme at the cell surface of WT, Δ*gtrR* and Δ*chdC* Mtb measured by pyridine hemochromogen. Data is shown as a ratio of ferrous to total heme and averaged from 2 independent trials. **d**. Effect of oxygen on exogenous heme bioavailability as measured by eGFP/mKATE2 ratio of HS1-M7A in WT Mtb in ambient air (+O_2_) or an anerobic chamber (-O_2_) for ∼18 hours. Cells are in PBS +/-5 µM Heme at 30 °C. For each independent experiment, ratio was normalized to the heme invariant sensor, HS1-M7A,H102A, to account for changes in fluorescence due to environment. To account for changes in fluorescence values between independent experiments, eGFP/mKATE2 ratios were normalized to Air (+O_2_, - Heme). Data shown is from 3 independent experiments. For (**b**) a one-way ANOVA with Dunnett’s post hoc comparison was used with WT as control, p = 0.0059.For (**c**) a one-way ANOVA with Dunnett’s post hoc comparison was used with WT as control. For Δ*gtrR* the open circle outlier was 1.6 standard deviations from the mean and was omitted from statistical analyses and ANOVA tests: ** *p* = 0.0092. For (**d**)) a one-way ANOVA with Tukey’s post hoc comparison was used, p = 0.0003. In all panels, ns denotes not significant.

This observed preference for transporting ZnPP over GaPP in WT Mtb led us to develop a <u>m</u>odified <u>p</u>yridine <u>h</u>emochromagen assay (MPH, see Supplemental methods) for determining the concentration of ferrous heme in Mtb cells. (38, 39) WT, Δ*gtrR* and Δ*chdC* Mtb cells treated with 25 µM heme for 7 days in 7H9 + ADS with aeration, all had ∼35-50% ferrous heme in whole cells lysed in <u>p</u>yridine <u>h</u>emochromagen <u>s</u>olution (PHS, see Supplemental methods) (**Fig. S9a**). The results in Δ*gtrR* and Δ*chdC* Mtb, which cannot synthesize heme, implied that ferric exogenous heme was reduced either during uptake, in the cytosol, or both (**Fig. S9a**). To assess whether Mtb could reduce extracellular heme, WT, Δ*gtrR* and Δ*chdC* Mtb cells were incubated with 5 µM ferric heme in PBS in an anaerobic chamber for 24 hours. Cells were pelleted and resuspended in PHS for 1-2 minutes and then pelleted again. Heme remaining in the cell-free PHS supernatant was measured by the MPH assay and is referred to as cell-surface associated heme, given that it was likely associated with the outer cell surface and/or possibly the periplasm. The MPH assay revealed that ∼1% of cell surface associated heme was reduced in WT Mtb (**Fig. 3c**). Most interestingly, Δ*chdC* cells exhibited a nearly 3-fold greater surface heme reductase activity than WT cells (**Fig. 3c**). Δ*gtrR* Mtb cells exhibited similar surface heme reductase activity to WT cells, suggesting that extracellular heme reduction is inhibited by ChdC and not general heme synthesis. In the presence of atmospheric oxygen (O_2_), we could not observe heme reduction at the cell surface, indicating that O_2_ could re-oxidize reduced heme. Disrupted cell lysates also contained a ferric heme reducing activity (**Fig. S9b-d**) that was lost upon boiling (**Fig. S9e**), indicating that the heme reductase activity likely arises from an enzyme (**Fig. S9b**,**c**). However, interestingly, unlike in intact cells, cell lysates from Δ*chdC* cells did not have an increase in heme reducing activity, suggesting that upon lysis, ChdC was decoupled from regulating heme reductase activity (**Fig. S9 d**,**f**).

The identification of heme reductase activity in Mtb led us to determine if exogenously supplied ferric heme is more bioavailable to cells in the absence of O_2_. Towards this end, we measured LH in WT Mtb expressing HS1-M7A after 24 hours in PBS supplemented with 5 µM exogenous heme in air versus an anaerobic chamber (**Fig. 3d**). Upon normalizing the sensor signal to the heme invariant HS1-M7A, H102A to account for heme-independent changes in fluorescence signal, which were minimal, we found heme bioavailability was significantly elevated in the absence of O_2_, consistent with a pathway for reductive heme uptake. Altogether, our results suggest that ChdC regulates heme import by inhibiting a reductive heme uptake pathway.

### Role of endogenous heme synthesis in macrophage infection

Given that Mtb may reside in macrophages where iron availability is limited to the pathogen,(40) we sought to determine if ChdC plays a role in Mtb survival and replication in the macrophage. Towards this end, we employed an *in vitro* assay of intracellular infection using RAW 264.7 macrophages in order to determine differences in metabolic activity between Δ*gtrR* and Δ*chdC* Mtb in response to macrophage infection (41). We measured the reduction of the tetrazolium dye MTT, which forms a purple formazan, as a readout of metabolic activity and viability in Mtb (42, 43). MTT-reducing activity of WT and heme depleted Δ*gtrR* and Δ*chdC* Mtb was measured before and 24-hours after RAW macrophage infection. Prior to infection, WT and heme depleted Δ*gtrR* and Δ*chdC* Mtb had similar metabolic activity (**Fig. 4a**). However, in Mtb cells isolated from macrophages after infection, both Δ*gtrR* and Δ*chdC* Mtb had a > 30 % decrease in metabolic activity compared to WT Mtb (**Fig. 4b**). This highlighted the importance of *de novo* heme synthesis during macrophage infection. There was also a difference in metabolic activity between the two heme auxotrophic strains, as Δ*chdC* Mtb cells had greater MTT-reducing activity than Δ*gtrR* Mtb cells. We suggest that this is due to the ability of Δ*chdC* Mtb to more efficiently acquire and utilize exogenous heme than Δ*gtrR* Mtb. However, we cannot rule out a role for heme biosynthetic pathway intermediates in macrophage survival. Altogether, our results indicated that both heme biosynthesis and uptake were important for survival in macrophages.

**Fig. 4.**
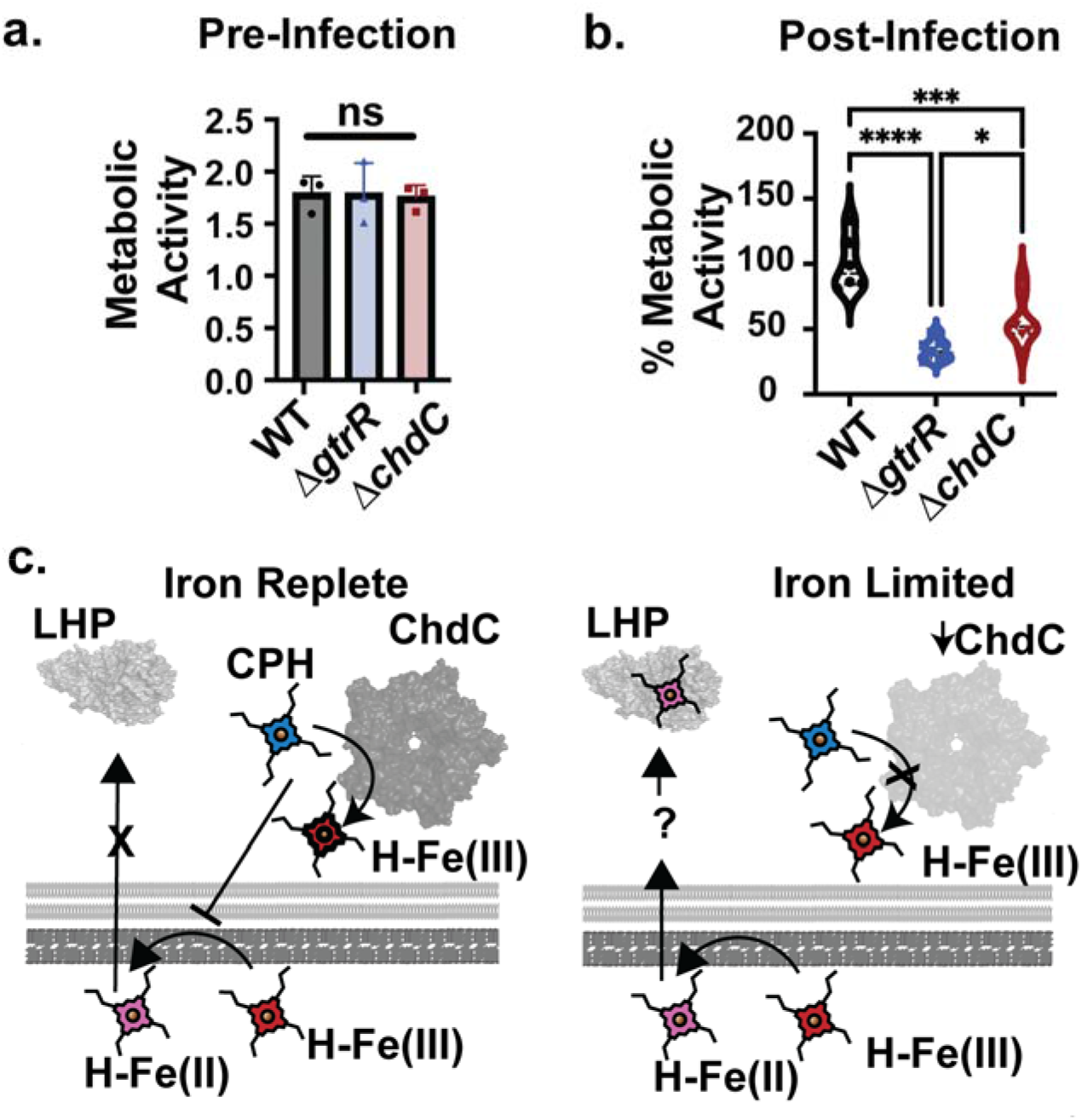
Role of heme synthesis in a macrophage infection model. **a**. Formazan absorbance as turnover of MTT in WT Mtb, Δ*gtrR* Mtb and Δ*chdC* Mtb of 2 × 10^6^ cells prior to macrophage infection. 2 × 10^6^ cells are equivalent to 100% engulfment and survival in macrophage infection. **b**. MTT absorbance of WT Mtb, Δ*gtrR* Mtb and ΔchdC Mtb isolated from RAW 246.7 macrophages after 24 hours of infection from 2 independent trials. WT Mtb n=6, Δ*chdC* Mtb n=8 and Δ*gtrR* Mtb n=7. **c**. Proposed model for the roles of ChdC in coordinating heme synthesis, uptake, and utilization. LHP = labile heme pool, CPH = iron coproheme, H-Fe(II) ferrous heme, H-Fe(III) = ferric heme. In panel **a**, data represent the mean ± S.D. (error bars) of triplicate cultures. The statistical significance was assessed by a one-way ANOVA with Dunnett’s post hoc test using WT Mtb as a control: “ns” denotes a non-significant *p* > 0.9689. In panel **b**, statistical significance was assessed by a one-way ANOVA with Tukey post hoc test: *** *p* = 0.0003 (WT vs. ChdCΔ); \**p* = 0.0423 (Δ*chdC* vs. Δ*gtrR*); **** *p* < 0.0001.

## Discussion

Heme is both an essential cofactor and a source of nutritional iron for the human pathogen Mtb (22). The importance of heme in Mtb physiology is underscored by the fact that there are at least two independent routes for heme acquisition, and heme can be synthesized *de novo* by Mtb (8, 10). To balance the nutritional and metabolic demands for iron and heme, while also mitigating the potential for heme toxicity, Mtb must tightly coordinate heme uptake, synthesis and utilization and integrate these processes with iron availability. Herein, our work identifies a previously uncharacterized mechanism in which the terminal heme biosynthetic enzyme, coproheme decarboxylase (ChdC), negatively regulates heme uptake and utilization through a reductive heme uptake pathway (**Fig. 4c**). Iron limitation down-regulates ChdC and increases heme import and bioavailability. (**Fig. 2 e-i**). We propose that such a mechanism may serve to protect cells from heme toxicity while trying to meet the nutritional demand for iron via heme scavenging and uptake from the host. These findings have a number of implications for iron and heme homeostasis in Mtb at the host-pathogen axis.

Heme is a cofactor necessary for Mtb virulence as heme dependent proteins such as KatG,(30, 44) cytochrome P450s,(45) and the Dos two component system,(46) are required for the survival and virulence of Mtb in infection models (23, 30). However, the relative contributions of *de novo* synthesized and exogenously scavenged heme towards labile bioactive heme pools and protein hemylation were unknown. Herein, we found that under conditions in which endogenously synthesized and exogenously supplied heme contributed equally to the total cellular heme concentration (**Fig. 1b**), exogenous heme was poorly utilized by mycobacteria and was inefficient at rescuing the growth of a heme deficient Δ*gtrR* strain (**Figs. 1a** and **2a**), contributing to the LH pool (**Fig. 1c**) and activating the heme enzyme KatG (**Fig. 1d)**. Additionally, using an Mtb-macrophage infection assay we found that survival of the heme synthetic mutants, Δ*gtrR* and Δ*chdC*, were impaired in infected macrophages compared to WT Mtb (**Fig. 4b**). Our findings that *de novo* synthesized heme is important for Mtb survival was also supported by prior studies indicating that many enzymes of the Mtb heme synthetic pathway were upregulated in Tb patient lungs compared with *in vitro* growth conditions.(47) We also found that LH levels were a good indicator of the bioactivity of subcellular heme pools in mycobacteria, including for activation of KatG, which is necessary for survival of Mtb in the macrophage(23, 30) and expressed in clinical isolates of Tb patients.(44)

One potential explanation for the apparent poor bioavailability of exogenously supplied heme is that it is degraded. However, we counterintuitively found that the transcript for the heme degrading heme oxygenase, MhuD, is repressed by heme supplementation and induced under heme deficient conditions (**Fig. 2d**), suggesting that heme degradation did not account for the differences in bioavailability between exogenously transported and endogenously made heme. Nonetheless, we cannot rule out this possibility as we have been unable to measure mycobilin, a product of heme degradation in mycobacteria, from cell extracts and therefore assess the extent to which heme is degraded *in vivo*.

An alternative explanation is that Mtb may regulate the trafficking and bioavailability of endogenously synthesized and exogenously scavenged heme differently due to differences in oxidation state and its concomitant effects on subcellular localization. Heme synthesized via ChdC is ferric, as ferric iron in coproheme is required for decarboxylation to heme.(48, 49) However, since the major mycobacterial redox buffer is mycothiol, which sets the cytosolic redox poise to highly reducing values, between -320 and -220 mV vs. the normal hydrogen electrode (NHE),(50) newly synthesized ferric heme may be rapidly reduced to ferrous heme, assuming its potential is similar to monomeric aqueous heme, (−50 mV vs. NHE),(1, 51) and labile heme equilibrates with the mycothiol redox buffer. Thus, one may reasonably expect that cytosolic heme is largely in its reduced ferrous state. In contrast, exogenous extracellular heme, which is in its oxidized ferric state, may gain access to the periplasmic space, which accounts for its contribution to “total” heme levels, but cannot be transported across the plasma membrane into the cytosol unless it is reduced by the ferric heme reductase system, thereby accounting for its comparatively low bioavailability. Ablation of ChdC, which otherwise serves to repress ferric heme reductase activity, results in elevated exogenous heme reduction (**Fig. 3c**) and uptake (**Figs. 2b** and **2c**), which would explain its increased availability in Δ*chdC* cells. Indeed, our findings that WT Mtb took up more ferrous heme analog, ZnPP, than ferric heme analog, GaPP, **(Fig. 3 a,b)**, and that ferric exogenous heme was more bioavailable in the absence of O_2_ (**Fig. 3d**) support the proposal that exogenously acquired heme is trafficked more efficiently into the cytosol upon reduction into its ferrous state. Therefore, in sum, heme iron oxidation state may be a key determinant that leads to the differential trafficking and bioavailability of exogenously scavenged and endogenously synthesized heme.

What is the nature of the ferric heme reductase and transport system? It likely requires electrons from NADH or NADPH and utilizes a flavin to ultimately reduce exogenous Fe^3+^-heme. Prior studies have implicated electron transfer flavoproteins, FixAB, as being required for utilization of heme as an iron source,(10) raising the intriguing possibility that FixAB are components of the ferric heme reductase system. The reductase system is likely present in the plasma membrane, where it would have access to cytosolically produced NAD(P)H. As such, we propose a model in which extracellular Fe^3+^-heme partitions into the periplasmic space, via outer membrane proteins such as porins or heme transporters, *e*.*g*. PPE62, or possibly diffusion through the cell envelope and outer-membrane, where it can be reduced by the ferric heme reductase system and transported across the plasma membrane into the cytosol.

Interestingly, deletion of ChdC disrupts the ability of Mtb cells to selectively uptake the ferrous heme mimic, ZnPP, over ferric heme mimic, GaPP (**Fig. 3b**). Given that both ZnPP and GaPP are redox inactive, it would suggest that ChdC is also required for the unknown heme transport system to confer selectivity for ferrous heme uptake, in addition to the ferric heme reductase activity. Taken together, we propose that when ChdC expression is increased in iron replete conditions, heme synthesis is active and heme uptake is repressed due to the inhibition of the ferric heme reductase and enhanced selectivity in transport of ferrous heme, thereby limiting uptake of exogenous oxidized heme. When ChdC expression is low, heme synthesis is diminished and heme uptake is elevated due to the de-repression of the ferric heme reductase and enhanced transport of both ferrous and ferric heme. Coupling heme reduction to ferrous selective transport using a heme biosynthetic enzyme may ensure Mtb does not inappropriately accumulate toxic levels of heme.

Exactly how heme reduction is coupled to its transport and the mechanisms underlying ChdC-mediated inhibition of heme reductase activity and selectivity in ferrous heme uptake remains to be determined. Since the ChdC-dependent heme reductase activity is lost in disrupted cell lysates (**Fig. S9b**), there may be key protein-protein interactions and a defined molecular architecture of ChdC and the heme reductase and transport systems akin to the heme biosynthetic metabolon discovered in the mitochondrial inner-membrane of eukaryotes.(52-54) Alternatively, since ablation of ChdC results in the inability to sense heme as evidenced by the lack of a transcriptional signature consistent with heme starvation in Δ*chdC* cells (**Fig. 2d**), ChdC may act as an intracellular sensor of heme that regulates the expression of additional factor(s) that control heme reductase activity and uptake.

Heme uptake pathways are important for Mtb survival in macrophage and mouse infection models.(8, 55) While iron levels are considered to be sufficient to support Mtb growth within the macrophage (56) or outer layers of the granuloma,(57) the caseous center of the granuloma is predicted to be severely iron limiting.(57) Our findings that iron limitation down-regulates ChdC protein level and increases heme import and bioavailability (**Fig. 2 g-i**) suggest that ChdC-regulated heme transport may be an important component of the iron starvation response in Mtb and is required for scavenging heme-iron from the host. While extracellular iron levels regulate the transcription of genes encoding other enzymes in the CPD pathway, including glutamate-semialdehyde amino mutase, GsaM, (also listed as HemL) and CpfC (also listed as HemZ),(57) the changes in *chdC* expression from multiple transcriptomic studies are distinct from other heme synthetic enzymes, further highlighting the unique role of ChdC in coordinating heme synthesis and uptake in response to iron availability. Further studies are needed to clarify the relationship between iron availability within the host and ChdC protein expression.

Iron regulation of ChdC likely occurs at the post-translational level given that prior transcriptomic studies have not shown that ChdC mRNA levels vary in response to changes in iron levels.(57, 58) ChdC is part of a subset of Mtb proteins (including only one other heme synthesis enzyme, GsaM), degraded via the CLP protease.(59) The CLP protease degrades proteins under non-stress conditions, suggesting ChdC is actively turned-over. Since ChdC catalytic activity is required to suppress heme uptake and utilization, it is tempting to speculate that iron regulation of ChdC expression occurs through a mechanism that involves differential turn-over of heme or coproheme bound and unbound forms of ChdC via the CLP protease.

The specific molecular mechanisms underlying ChdC-mediated regulation of heme uptake, bioavailability and utilization also remains to be determined. While we have shown that ChdC regulates a reductive heme uptake pathway, we were unable to transcriptionally identify known heme transport genes involved in ChdC regulated heme uptake (**Fig. 2d**). We predict that as-yet-to-be determined ChdC protein-protein interactions are involved in regulating factors that control heme reduction, uptake and bioavailability. For example, in *Staphylococcus aureus*, ChdC interacts with coproporphyrin ferrochelatase (CpfC) and IsdG, one of two heme degrading enzymes.(60, 61) Given that catalytically active ChdC is required to suppress heme uptake and bioavailability (**Fig. S4**), we propose that transfer of heme from ChdC to a downstream target inhibits heme reduction, uptake and/or bioavailability. For instance, protein-protein interactions have been identified that transport newly synthesized heme from the terminal enzyme of the PPD branch, ferrochelatase, to downstream hemoproteins.(62) Alternatively, it is possible that heme-bound ChdC has a distinct interaction network from heme-free ChdC to control heme uptake. Identification of the ChdC interactome will clarify how ChdC regulates heme uptake and utilization.

Post translational modifications (PTMs) of ChdC may be critical for regulating its expression and/or interactions with factors that regulate heme uptake and bioavailability. Mtb ChdC is both succinylated (63) and pupylated. (64) Both succinylation and pupylation occur at surface exposed lysines and could inhibit protein-protein interactions, or in the case of pupylation, also mark the protein for degradation.(64) Interestingly, pupylation has already been shown to be an iron dependent regulator of ferritin in *Corynebacterium glutamicum*.*(65)* In *C. glutamicum*, ChdC is also pupylated, however, no pup-dependent proteasomal machinery exists.(66) Moreover, current studies do not support pupylation dependent degradation of ChdC, suggesting pupylation of ChdC may play a regulatory role.(67) Further studies are warranted to clarify how ChdC PTMs affect expression, activity, and regulation of heme uptake and bioavailability.

One surprising finding from our work was the level of free porphyrins in WT Mtb (**Fig. S8)**. The buildup of heme intermediates is considered disadvantageous due to the inherent cytotoxicity of porphyrins. In Msm, the build-up of porphyrins in cultures during the transition to dormancy increases the susceptibility of Msm to photoinactivation. (68) As has been proposed for the treatment of other bacteria(69) and cancers, the exploitation of light enhanced porphyrin toxicity may underly the basis of future anti-Mtb therapies. (70)

Altogether, our studies are the first to demonstrate that the regulation of heme synthesis, uptake and bioavailability in mycobacteria requires the heme synthetic enzyme, ChdC, whose expression is both iron and heme regulated, and which controls a reductive heme uptake pathway (**Fig. 4c**). A detailed knowledge of how ChdC coordinates heme uptake and utilization and the specific biological contexts in which it does so will be essential for understanding how Mtb adapts to varying availability of iron, heme and oxygen during infection.

## Materials and Methods

All materials, including reagents, cell lines, culture conditions, and methods are described in detail in the *Supplemental Material and Methods*. Mtb strains used in this work are avirulent and in the mc^2^6230 (H37Rv Δ*rd1* Δ*panCD*) background. Msm strains are in the mc^2^155 background. Generation of knockout strains and sensor incorporation in mycobacteria are also described in *Supplemental Material and Methods*. All experimental methods including total heme assay,(71) labile heme,(16) sensor calibration,(16) porphyrin assay,(71) catalase-peroxidase activity assay, (24, 25) fluorescent porphyrin assays, and ferrous heme measurements are also described in *Supplemental Material and Methods* in the Supplemental Appendix.

## Acknowledgment

We thank Prof. William R. Jacobs, Jr. (Albert Einstein College of Medicine) for sharing *M. smegmatis* mc^2^155, *M. tuberculosis* mc^2^6230, and the specialized transduction system and base plasmids used in this work. We thank Prof. Daniel Kornitzer for helpful discussions regarding heme reductase activity. We wish to acknowledge the core facilities at the Parker H. Petit Institute for Bioengineering and Bioscience at the Georgia Institute of Technology for the use of their shared equipment, services and expertise.

