## Supporting Information and Figures for "The terminal heme synthetic enzyme, Coproheme Decarboxylase, coordinates heme synthesis and uptake in response to iron in *Mycobacteria*"

Paras Jain

Trudeau Institute

154 Algonquin Ave,

Saranac Lake, NY 12983

**This PDF file includes:**

Supplementary Text

Figures S1 to S9

Tables S1 to S2

SI References

Supplemental Results

**Role of ChdC catalytic activity in heme uptake and utilization.** We sought to determine if enzymatic activity was necessary for ChdC to regulate heme uptake and utilization in Msm as assessed by KatG activity. Towards this end, we expressed a series of Mtb *chdC* alleles with mutations in the predicted active site in ∆*chdC* Msm **(Fig. S4a-c).** The H156A mutant of Mtb ChdC was previously shown to be unable to bind heme and to have reduced catalytic activity as iron coordination by histidine was necessary for catalysis.(1) The Y133S/R137A mutant had been shown to have reduced catalytic activity and abrogated coproheme binding,(2) and the H156A/Y133S/R137A mutant was expected to be defective in both catalysis and heme binding. WT Mtb ChdC complemented ∆*chdC* Msm as expression of WT Mtb *chdC* rescued ∆*chdC* Msm in the absence of added heme (**Fig. S4d)** In contrast, the aforementioned ChdC mutants did not restore growth (**Fig. S4e**) in ∆*chdC* Msm cells not supplemented with heme. Similar to growth, Mtb ChdC restored KatG activity in the absence of heme, however all three ChdC variants had similar KatG activity to ∆*chdC* Msm (**Figs.** **S4f,g**). These results indicated that the three ChdC mutants were catalytically inactive. As with ∆*chdC* Msm, and unlike ∆*gtrR* cells, ∆*chdC* cells expressing catalytically inactive ChdC supplemented with exogenous heme exhibited increased KatG activity and grew similarly to *∆chdC* Msm (**Figs. 3g**, **S4e,f**). Taken together, these data suggested that the enzymatic activity of ChdC, and not just the presence of the ChdC polypeptide, was required to negatively regulate heme uptake and utilization.

**Exogenous heme decreases porphyrin synthesis in *M. tuberculosis.*** Heme synthesis is regulated by intracellular heme levels in some bacteria.(3) To determine if exogenous heme could feedback to regulate heme synthesis in mycobacteria, we measured fluorescent porphyrins as a proxy for flux through the heme biosynthetic pathway. Porphyrins fluoresce at ~ 650 nm when excited with 400 nm light. The levels of these species, which we refer to as “free porphyrins” (FPs), would include the heme biosynthetic intermediate coproporphyrin along with any porphyrinogens that oxidize in air upon cell lysis. An increase in FPs would indicate that there is either a block in late stages of heme synthesis or that iron insertion by ferrochelatase could not keep up with increased flux through the heme synthetic pathway.

We found that WT Mtb lysates exhibited a steady-state level of porphyrin fluorescence emission (**Fig. S8a,b**). When these cells were treated with the heme synthetic inhibitor, SA, FP emission decreased, consistent with the SA-mediated inhibition of porphobilinogen synthase, an early step of heme synthesis (**Fig. S8a,b**). In support of this, the 650 nm emission peak was greatly reduced in ∆*gtrR* Mtb and Msm strains (**Fig. S8 c,d**), but increased in ∆*gtrR* Msm supplemented with ALA, due to increased porphyrin synthesis (**Fig. S8c)**. Most notably, heme addition to WT Mtb decreased FP emission to levels similar to those measured with addition of SA, which indicated that heme acted as a feedback inhibitor of heme synthesis (**Fig. S8ab**). We also observed that exogenous heme suppressed porphyrin accumulation in ∆*chdC* Mtb and Msm. Both ∆*chdC* Mtb and ∆*chdC* Msm exhibited increased FP emission because of the block in the last step of heme synthesis (**Fig S8 c,f**). The increased FP emission of ∆*chdC* strains was reversed by addition of exogenous heme, which further indicated that heme inhibited porphyrin synthesis (**Fig. S8 c,f**). Finally, FP levels in WT Mtb were increased by addition of ALA but decreased with exogenous SA and Hb (**Fig. S8g**). Interestingly, heme dependent inhibition of FP accumulation was not observed in WT Msm (**Fig. S8h**). Taken together, our data indicated that exogenous heme inhibited heme synthesis in WT Mtb.

**Supplemental Materials and Methods**

**Bacterial strains and reagents.** The *M. smegmatis* mc^2^155 and *M. tuberculosis* mc^2^6230 (H37Rv *Δrd1* *ΔpanCD*) were obtained from laboratory stocks. For knockouts and transformations, the strains were grown in Middlebrook 7H9 (Difco, Sparks, MD) supplemented with 10% (vol/vol) oleic acid-albumin-dextrose catalase (OADC; Difco), 0.2% (vol/vol) glycerol, and 0.05% (vol/vol) tween80 at 37^o^C with shaking. Middlebrook 7H10 (Difco) supplemented with 10% (vol/vol) OADC and 0.2% (vol/vol) glycerol was used as solid medium. The following supplements were used, as necessary, at these concentrations: L-pantothenate 50 µg/ml; ALA 1.5 µg/mL; heme 25 µM. The plasmid pYUB1471, shuttle phasmid phAE159, and phage phAE280 were obtained from laboratory stocks (4). Hygromycin (Gold Biotechnology, St. Louis, MO) was used at concentrations of 50 µg/ml for mycobacteria and 150 µg/ml for *Escherichia coli***.** All the supplements were obtained from Sigma-Aldrich or Thermo (Fisher) Scientific. All primers used for generation of mutant and sensor strains are listed in **Supplemental Table 1.**

**Generation of ∆*gtrR* and ∆*chdC* mutants.** The genes for *gtrR* and *chdC* were identified by bioinformatics analysis in *M. smegmatis* and *M. tuberculosis*. The deletion mutants of *gtrR* and c*hdC* were generated by specialized transduction as described previously.(4) The transductants were selected on plates containing hygromycin as the selective marker and either ALA or heme. The hygromycin cassette was excised from the knockout strains using the phage phAE280 and sucrose selection (4). The deletion and unmarked strains were confirmed by PCR and sequencing. The primers used to generate and confirm the mutant strains are listed in SI.

**Generation of heme sensors for mycobacteria.** The heme sensors HS1, HS1-M7A and HS1-M7A-H102C were cloned under the control of G13 promoter in an episomal mycobacterial plasmid (5-7). The resulting plasmids expressing HS1 (pYUB1872), HS1-M7A (pYUB1874) and HS1-M7A-H102C (pYUB1876) were confirmed by restriction digestion and sequencing. Plasmids were electroporated in various strains of mycobacteria using the following settings (2.5 kV, 25 mF, and 1,000 ohms) (8). Transformants were selected on 7H10 plates containing necessary supplements and Kanamycin 40 µg/ml. Plates were incubated at 37°C for 3 days for Msm and 4 to 6 weeks for Mtb.

**Generation of ChdC mutants for complementation.** Plasmids expressing Mtb *chdC* alleles were constructed using standard PCR cloning methods. Open reading frame (ORF) encoding HS1 in pYUB1872 was excised NdeI and NheI restriction enzymes and replaced by Rv2676c ORF using NEBuilder HiFi DNA Assembly. The resulting plasmid (pTIP19) constitutively expressed wild type Mtb *chdC* from G13 promoter. Plasmids expressing Mtb *chdC* mutant alleles ChdC Y133S/R137A (pTIP20), ChdC H156A (pTIP22) and ChdC H156A/Y133S/R137A (pTIP24) were generated by site directed mutagenesis of pTIP19 using mutation primers and NEBuilder HiFi DNA Assembly. Successful incorporation of mutation/s was confirmed by sequencing. The primers used are listed in S1. Plasmids were transformed in Msm strains as described above.

***M. smegmatis* growth and heme depletion.** Msm strains were grown shaking at 170-200 rpm at 37 °C in Middlebrook 7H9 media (Difco) supplemented with 10% ADS (albumin dextrose and salt) and 0.05 % tween 80. ADS was composed of 5% bovine serum albumin (Fraction V Gemini Biosciences), 2% Glucose and 0.85% sodium chloride in water. Sensor strains were grown with 40 µg/mL Kanamycin sulfate. ∆*gtrR* and ∆*chdC* Msm strains were supplemented with 1.5 µg/mL ALA or 25 µM hemin chloride respectively. For depletion, cells were grown in ALA or heme supplemented media to an OD_600_ of ~ 1. Cells were pelleted and washed 3x with sterile water before resuspension at 1 OD_600_ in 7H9 media with no ALA or heme. Cells were grown in depleted media for 18 hours. After depletion, cells were resuspended at 1 OD_600_ and measurements taken as heme depleted cells. Cells were then supplemented with 5 µg/mL ALA or 50 µM hemin chloride for ALA and heme studies.

***M. tuberculosis* growth and heme depletion.** Mtb cells were grown shaking at 170-200 rpm at 37 °C in Middlebrook 7H9 media (Difco) supplemented with 10% ADS, 0.2% Casamino acids (Difco), 0.02% Tyloxapol, and 50 µg/mL calcium pantothenate (Sigma). Cells were grown in 10-11 mL cultures in closed 30 mL square PETG bottles (Fisher). ∆*gtrR* strains were grown with 5 µg/mL ALA and ∆*chdC* strains were grown with 25 µM hemin chloride. For heme depletion, ∆*gtrR* and ∆*chdC* strains cells were pelleted, washed 1X in 7H9 media and resuspended in 7H9 media without hemin or ALA supplementation. Sensor strains were grown with 40 µg/mL kanamycin.

***M. tuberculosis* growth in minimal media.** Mtb were grown from freezer stocks in 7H9 as above until they reached saturation (~ 10 days for WT and 20-30 days for ∆*gtrR* and ∆*chdC* strains). Cells were washed 1x with minimal media not supplemented with Fe, then diluted into minimal media. Minimal media was a modified M63 minimal media containing 1X M63 salts (no Fe added) with 2% glucose, 0.5% casamino acids, and 50 µg/mL calcium pantothenate was added. Fe was added as ferrous ammonium sulfate dissolved in water. Heme was added as hemin chloride solubilized in 100% DMSO. Sensor strains were grown with 40 µg/mL kanamycin.

**Labile heme assay**. Cells at each time point were taken, and the OD_600_ measured to determine cell density. Cells were pelleted and washed in water 2X before being suspended in phosphate buffered saline (PBS) at an OD_600_ of 1 for Mtb and of 10 for Msm for sensor fluorescence measurements. For sensor fluorescence, 200 µLs of cells were plated in 96 well flat bottom Grenier Fluorotrac plates as technical duplicates. WT Msm and Mtb cells not expressing sensor were measured and subtracted as background fluorescence for both the eGFP and mKATE2 channels. Sensor fluorescence was measured on a Biotek Synergy MX plate reader, with an excitation at 480 nm and emission at 510 nm for eGFP, and an excitation at 580 nm and emission at 620 nm for mKATE2, slit widths for excitation and emission were 9 nm. Multiple reads over 5-10 minutes were taken to account for variability in fluorescence over time. Multiple reads and technical replicates were averaged as one ratio. The reported mean and standard deviation are calculated from biological replicates.

***In situ*** **heme sensor calibration.** In order to relate the sensor fluorescence ratio to the fractional heme occupancy of the sensor, a previously established method to calibrate the heme sensors in yeast was adapted for mycobacteria.(5)

$\% Bound =\left( \frac{R_{expt}-R_{min}}{R_{max}-R_{min}} \right)x 100$ (**Equation 1**)

The amount of heme bound to the sensor, % Bound, can be quantified by determining the sensor eGFP/mKATE2 fluorescence ratio under any given experimental condition, *R*_expt_, relative to the eGFP/mKATE2 fluorescence ratios when the sensor is 0% (*R*_min_) and 100% (*R*_max_) bound to heme, respectively. The theoretical limit for *R*_max_ is ~ 0 due to the > 99% efficiency of energy transfer between GFP and heme. This was confirmed by permeabilizing cells and adding 50 μM hemin chloride, which saturated HS1 and gave a ratio approaching 0 (**Fig. S1d)**. *R*_min_ was determined by growing parallel cultures of cells expressing HS1-M7A,H102A, which cannot bind heme (**Fig. S1f)**.

To calibrate the sensors, cells were washed with water and resuspended in PBS at an OD_600_ of 10 and pre-permeabilization fluorescence was measured as detailed in the LH assay above. Cells were resuspended in a previously reported permeabilization buffer used for TUNEL assays in Mtb.(9) The permeabilization buffer of 0.1% Triton X-100 in 0.1% sodium citrate was supplemented with 1 mM ascorbate. To saturate the sensor, cells were treated with and without 50 µM hemin chloride for 30 minutes while shaking at 37 ˚C. Cells were pelleted, washed and resuspended in 1mM ascorbate in PBS and sensor fluorescence was measured and reported as permeabilized (no heme) and saturated (plus heme).

**Total heme assay.** Total heme fluorescence was based on a previously reported porphyrin fluorescence assay.(10) Briefly, Msm or Mtb cells were taken at time points and washed as for labile heme measurements above. After suspension at an OD_600_ of 10 in PBS form Msm or ~1 OD_600_ for Mtb, 500 µLs of cells were pelleted and frozen at -80 °C. Cells pellets were resuspended in 500 µLs of 20 mM oxalic Acid and allowed to sit at 4 °C overnight. To each cell suspension, 500 µLs of 2M oxalic acid were added. Cells were mixed and divided into two 500 µL aliquots. One was stored at in the dark at room temperature as a blank. The other was boiled at 100 ^0^C for 30 minutes covered. Blanks and boiled samples were centrifuged at 21,100 x g for 2 minutes to remove cell debris. For fluorescence measurements, 200 µLs of supernatant were plated as technical duplicates in 96 well black flat bottom Grenier Fluorotrac plates. Fluorescence was measured on a Tecan Infinite 200 Pro plate reader with excitation at 400 nm and emission at 608 nm and 662 nm. Emission spectra from 600-700 nm were also recorded. Heme concentration was calculated from fluorescence by comparison with heme standards from 1 nM to 500 nM treated with oxalic acid as cell samples above.

**Free porphyrin assay**. Free porphyrin measurements were based on a previously reported assay (10). Briefly, Msm and Mtb cells from the total heme assay were used. As in the total heme assay, cells were treated with 20 mM oxalic acid overnight. 2 M oxalic acid was added to samples and they were allowed to sit for 30 minutes in the dark. Free porphyrin was measured as fluorescence on a Tecan Infinite 200 Pro plate reader with excitation at 400 nm and emission from 600-700 nm. Cells are measured at equivalent OD_600_ and values are reported in AFU.

**Catalase-peroxidase activity gel** Catalase-peroxidase activity gels were based on a previously reported assay (11, 12). Briefly, cells were washed as with labile and total heme measurements. Cells at 10 ODs in PBS were collected, pelleted and frozen at -80 °C. Cells were resuspend in lysis buffer, PBS + 0.1 % Triton X-100 1mM EDTA and 1X protease arrest, and lysed with 0.5 mm zirconium oxide beads in a Bullet Blender tissue homogenizer at setting 8 for 3 minutes at 4 °C. Cell debris and beads were removed by centrifugation at 21,100 x g for 5 minutes. Lysate was measured for total protein content via Bradford or absorbance at 280nm. Even loading of total protein was verified by SDS PAGE. For catalase gels, lysates were separated on a 14% native Novex precast Tris-glycine gel at 4 °C for 16-18 hours. The catalase gel was washed 3 x 15 minutes in water then incubated with 0.3% H_2_O_2_ for 10 minutes. The gel was rinsed and added to a mixture of equivalent volumes (~30 mLs total) of 2% potassium ferricyanide and 2% iron chloride to stain the gel. The gel was rocked by hand until a faint green color started to appear in the gel, then the gel was transferred to water and imaged. Catalase bands appear clear or yellow on the green background of the gel. Gel images were converted to grey scale and inverted for easier viewing in print.

**Zinc mesoporphyrin uptake.** Msm and Mtb cells were treated with or without 1 µM ZnMP in culture in 7H9 media or MM for 10 hours for Msm, and 2, 24 or 48 hours for Mtb. ZnMP from Frontier Scientific was used from a 5 mM stock in DMSO store at -20 ˚C. Cells were washed 2x in PBS with 2% BSA to remove free ZnMP and then washed 2x in PBS. Cells were resuspended in PBS and lysed with 0.5 mm zirconium oxide beads (Msm) or 0.1 mm glass beads (Mtb) in a Bullet Blender tissue homogenizer at setting 8 for 3 minutes at 4 °C. Lysed cells were allowed to rest for 3-5 minutes before a second round of lysis at setting 8 for 3 minutes. Total protein was measured by absorbance at 280 nm to ensure even loading of lysates. 200 µLs lysate with equivalent total protein was added to a 96 well black flat bottom Grenier Fluorotrac plates. Fluorescence was measured on either a Tecan Infinite 200 Pro plate reader or a Biotek Synergy MX plate reader with excitation at 418 nm and emission spectra from 550-700 nm were recorded. Fluorescence spectra were baseline subtracted.

**RT-qPCR.** Cultures (10ml) of WT Mtb, Mtb ΔgtrR, Mtb ΔchdC were grown in triplicate under various test conditions. The cultures were centrifuged, and pelleted cells were resuspended in 1 mL RNA later stabilization solution overnight at 4 ˚C before storage at −80°C. For RNA extraction, the cells were centrifuged and resuspended in 1ml of TRIzol reagent (ambion). The samples were transferred to 2ml screwcap tubes containing 0.5mm Zirconia beads and processed in PowerLyzer 24 homogenizer (Qiagen) for 45s, 3500rpm (x4). The debris was spun down, and the supernatant (∼750 µL) was transferred to the fresh tube. Equal volume of absolute ethanol was added and applied to Zymo-Spin IIICG column. The RNA was extracted by following Direct-zol RNA miniprep plus protocol which includes on-column DNase digestion step. RNA yields were determined using Qubit RNA BR assay (Invitrogen). cDNA was synthesized using ∼400ng RNA and LunaScript RT supermix (NEB) in a 25 µL reaction using the following program: 2 min at 25°C, 20min at 55°C, 1 min at 95°C . No-RT control mix (NEB) was used to eliminate DNA contamination in samples. The cDNA products were quantified in triplicate by real-time PCR, using Luna universal qPCR mastermix (NEB) in a 20 µL reaction and 0.25 µM primer concentrations, and running the following program on an ABI 7500 fast real time system. 2 min at 95°C, 45 cycles of 15 s at 95°C, 30 s at 60°C (+plate read, SYBR), followed by a dissociation stage of 15 s at 95°C, 15 s at 60°C and 15 s at 95°C to check specificity of the products. Threshold cycles were normalized to those for 16S rRNA. Representative samples were also run on a 1.5% agarose gel. All primers used for qPCR are listed in **Supplemental Table 2**.

**Western blots.** For Western blots, cells were adjusted to the same OD_600_ before lysis and lysed in PBS with 1% SDS, 1mM EDTA, 0.1% Triton X-100 and 1X Protease Arrest (G Biosciences). Protein was loaded on a precast 14% gel and transferred to a 0.45 µM nitrocellulose membrane. For Ponceau staining, membrane was washed 2X in deionized water and stained with Ponceau S (VWR) dye for ~ 10 minutes. Blots were washed with water to remove Ponceau background and imaged. After imaging, blots were washed in TBS-T until all red color was removed. Blots were blocked with Licor Odyssey buffer. Blots were probed via a ChdC antibody (1:500) (provided by Dailey lab UGA) raised in rabbit and Alexa Fluor 680 goat anti-rabbit secondary antibody (1:1000). Blots were imaged using a Licor Odyssey imager. Ponceau S staining and ChdC expression was quantified using Li-Cor ImageStudio^TM^ Lite software.

**GaPP and ZnPP uptake.** Gallium(III)protoporphyrin IX (GaPP) and zinc(II)protoporphyrin IX (ZnPP) were both purchased from Frontier Scientific. GaPP and ZnMP stocks were made by dissolving GaPP and ZnPP in DMSO. Concentrations were measured by extinction coefficients reported in the literature.(13) GaPP and ZnPP at 1 µM or an equivalent volume of DMSO as control, were added for time shown or 48 hours in media (7H9+ADS) or BSA as described. Cells were placed at same OD, washed in 2 times in 2% BSA in PBS and i2 time in PBS to remove unbound GaPP or ZnPP. Fluorescence spectra of cells were measured with an excitation of 410 nm (9 nm badwidth) and emission spectra were read from 500 – 700 nm in 1 nm steps. With peak emission at 585 nm read for GaPP and 589 nm for ZnPP. Fluorescence in DMSO treated samples was subtracted and concentration per cell was calculated from a standard curve to account for different quantum yields for GaPP and ZnPP.

**Modified pyridine hemochromagen assay.** Ferrous heme and %ferrous heme were measured by an assay modified from the standard pyridine hemochromagen assay.(14) Briefly, cell pellets or lysates were degassed by 3 vacuum and gas cycles in an airlock and put in a COY brand anerobic chamber at 30 ˚C with an atmosphere of N_2_ and H_2_. Cell pellets were resuspended in degassed PBS. A heme stock in DMSO was degassed and diluted in degassed PBS. 5 µM heme was added to samples or degassed PBS as a control and incubated overnight. For the pyridine hemochromagen assay, the next day fresh pyridine hemochromagen solution (PHS) of 40% pyridine in 0.2 M NaOH (without potassium ferricyanide) was made in a chemical hood. PHS was aliquoted at 500 µLs – 1 mL, sealed and placed in the anaerobic chamber. To decrease dissolved oxygen, PHS caps were opened periodically over 1-2 hours. For cell surface reduced heme measurements, cells were pelleted, and supernatant removed, cells were resuspended in 500 µLs PHS for 1-2 minutes then pelleted. 400 µLs PHS supernatant was added to 400 µLs PBS in a quartz tube and sealed with a rubber cap to limit exposure to O_2_. Absorbance spectra from 500 – 600 nm was measured in an Olis Clarity UV-Vis. Two spectra were averaged for each sample. To measure total heme, 10 µLs of 0.5 M sodium dithionite in 0.5 M NaOH was added to completely reduce heme. Spectra were blank subtracted with 400 uLs of 5 µM heme in PBS that had incubated in the COY chamber overnight and baseline corrected to account for baseline drift. An extinction coefficient of 34.7 mM^-1^ cm^-1^ at 557nm was used as a measure of ferrous heme.(14)

**Macrophage infection assays.** Raw 246.7 macrophages were supplied at passage 4 from the Wood lab (Georgia Tech). Cells were grown in DMEM plus glutamine without antibiotics. Cells used for this assay were between passage 6 and 10. Cells were grown to 100% confluency in Grenier CellStar Tissue Culture Treated 6 well plates. Macrophage cells were washed with sterile Dulbecco’s PBS (DPBS) and fresh media was added prior to infection with Mtb. WT Mtb, ∆*gtrR* Mtb and ∆*chdC* Mtb were grown from freezer stocks to an OD_600_ ~ 1 as described above. WT Mtb and ∆*chdC* Mtb were diluted 1:10 into fresh 7H9/ADS media with 25 µM hemin chloride and ∆*gtrR* Mtb was diluted into 7H9/ADS media with 25 µM hemin chloride and 5 µg/mL ALA and Mtb cells were grown ~ 7 days until OD_600_ ~ 1. Mtb cells were depleted of heme for 2 days by washing with 7H9 media and resuspending in fresh 7H9/ ADS without hemin chloride. For infection Mtb cells were washed and resuspended in sterile DPBS at an OD_600_ of 4 (100X). Mtb cells were added to macrophage cells at an MOI of 10 (OD_600_ of 0.04) for 3 hours incubated at 37 °C in 5% CO_2_. 3 wells of macrophage treated with PBS were used as control. After 3 hours, extracellular Mtb was washed away with three washes with sterile DPBS and fresh DMEM was added with 40 µg/mL Kanamycin to kill any extracellular Mtb. Cells were incubated for 24 hours at 37 °C in 5% CO_2_. Cells were washed with sterile DPBS and lysed with 500 µLs sterile filtered 0.05% SDS in DPBS for 5 min at room temperature. To aid in lysis, SDS was pipetted up and down. 400 µLs of lysate was pelleted, washed with PBS and resuspended in 200 µLs PBS. 100 µLs of each sample was added in duplicate to a 96 well plate along with 20 µLs sterile filtered 5 mg/mL MTT in water. The MTT reaction was incubated at 37 °C for ~20 hours. Empty wells in plates were filled with sterile water and plates were sealed with parafilm to decrease evaporation. After 20 hours, 100 µLs 10% SDS in water was added to dissolve formazan crystals. This mixture was placed at 37 °C for four hours and then absorbance was read at 570 nm and 690 nm. The 690 nm absorbance was subtracted as background. Readings for macrophage only wells were greater than 5-fold lower than any Mtb infected well readings. These values were subtracted as background from wells infected with Mtb. For the Mtb pre-infection MTT assay, after cells were resuspended at an OD_600_ of 4 in sterile DPBS 16 µLs of cells were diluted to 200 µLs in sterile DPBS and added in duplicate 2 x 100 µLs to a 96 well plate. 20 µLs sterile filtered 5 mg/mL MTT was added and cells were incubated 37 °C for 20 hours, SDS added for 4 and then absorbance read all as above in macrophage treated cells.

**Supplemental Figures**

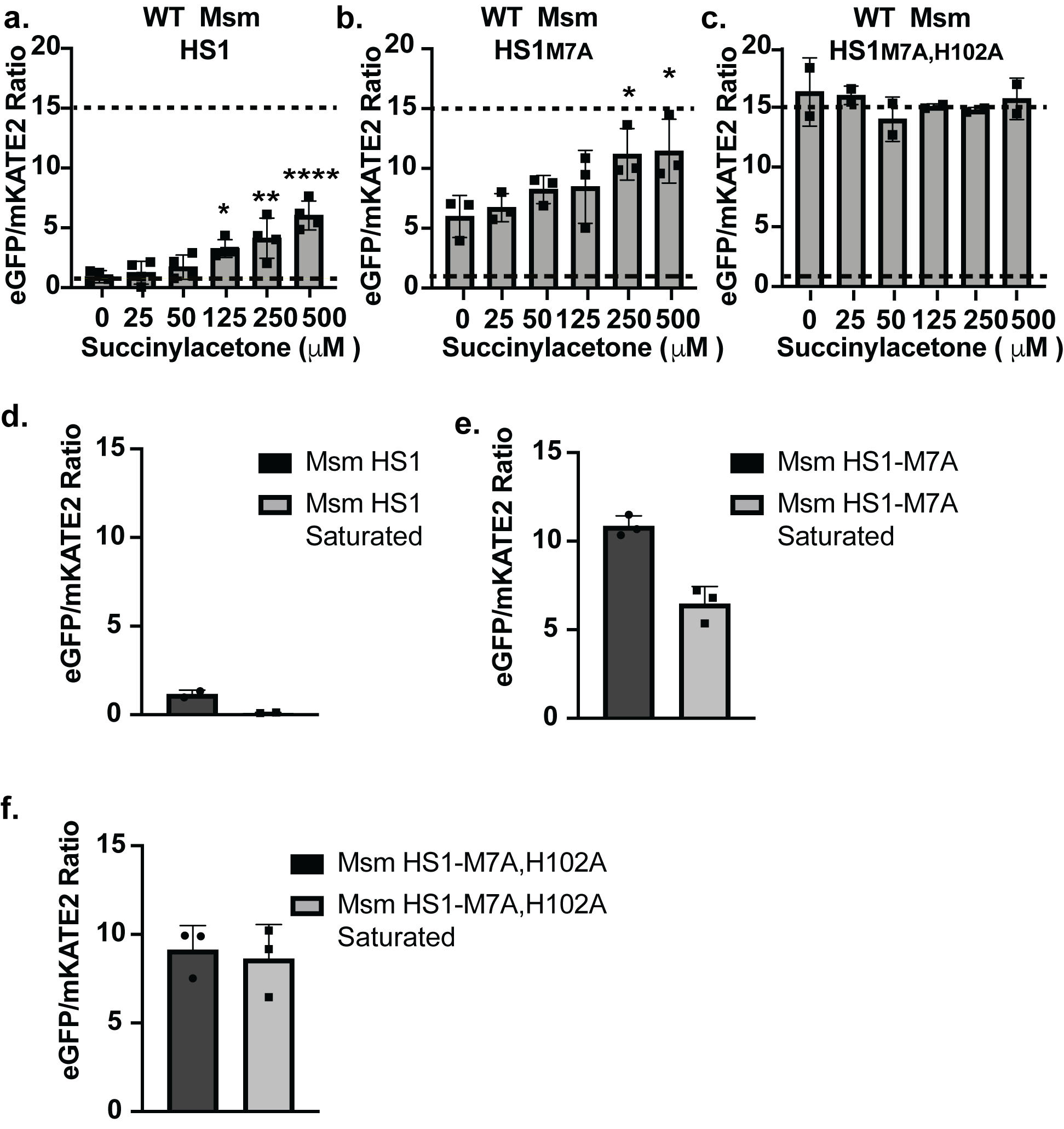

**Supplemental Figure 1. Validation of heme dependent response in Msm of HS1 and HS1-M7A and *in situ* LH sensor saturation.** The eGFP/mKATE2 fluorescence ratio of (**a**) HS1, (**b**) HS1-M7A, (**c**) and HS1-M7A,H102A expressed in WT Msm as a function of succinylacetone (SA), a heme synthesis inhibitor. **d.** WT Msm HS1, **e.** WT Msm HS1-M7A and **f.** WT Msm HS1-M7A,H102A before and after permeabilization with sodium citrate and Triton X-100 and saturation with 50 µM hemin. Data in all panels represent the mean ± S.D. (error bars) for n=3 except for panel **d.** where n=2. In panels **a-c,** the statistical significance was assessed by one-way ANOVA with Dunnett’s post hoc test using untreated Msm (0 µM SA) as the reference: (**a**) * *p* = 0.0264, ** *p* = 0.0023, *****p* = < 0.0001; (**b**) * *p* = 0.0417 and 0.0316.

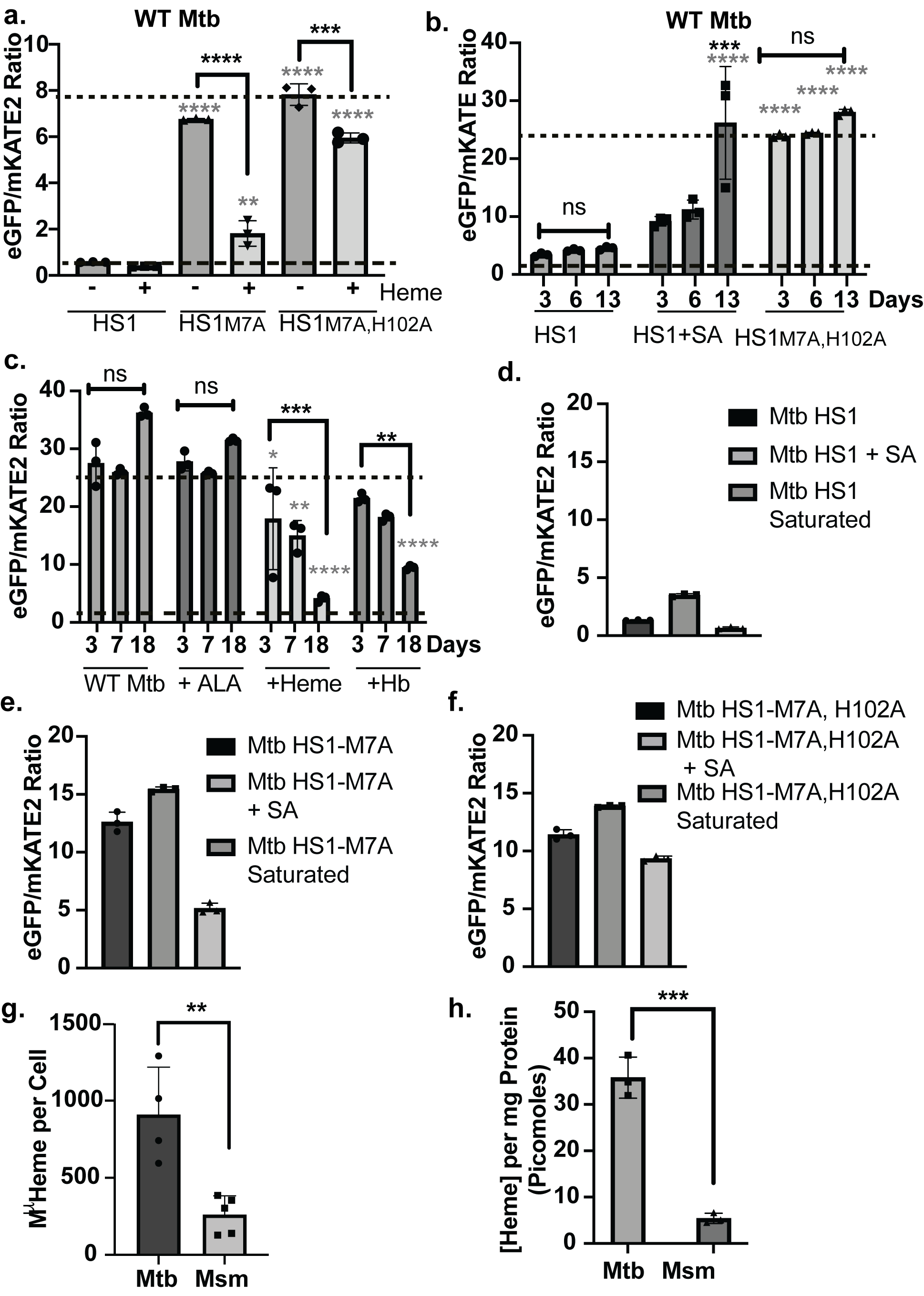

**Supplemental Figure 2. Validation of heme dependent response in Mtb of HS1 and HS1-M7A and *in situ* LH sensor saturation. a**. Fluorescence ratios of the indicated heme sensors expressed in WT Mtb cells conditioned with or without 25 μM hemin chloride for 13 days. **b**. Fluorescence ratio of HS1 expressed in WT Mtb cells conditioned with or without 500 μM SA over time. **c**. Time-dependent fluorescence ratio of HS1-M7A expressed in WT Mtb cells conditioned with 5 µg/mL of the heme precursor 5-aminolevulinic acid (ALA), 25 µM hemin chloride or 6.25 µM hemoglobin (Hb) . In panels **a-c**, fluorescence ratios indicative of 0% and 100% heme bound to sensor as determined by sensor calibration experiments is denoted by small and large dashed lines, respectively. **d.** WT Mtb HS1 **e.** WT Mtb HS1-M7A, and **f.** WT Mtb HS1-M7A,H102A with no treatment, treatment with 500 µM SA or after permeabilization with sodium citrate and Triton X-100 and saturation with 50 µM hemin. **g-h.** Concentration of total heme in WT Msm and WT Mtb cells (**g**) and lysates (**h**). (**g)**. Cell number was measured by OD_600_. Previously published cell volumes of Mtb and Msm were used.(15) Mtb n=4 and Msm n=5. Error shown is S.D. **P-value = 0.0033 from unpaired two-tailed T-test. All error bars shown are S.D. of n=3 in (**a-f**) and in (**h)**. In panel **a**, the statistical significance was assessed by one-way ANOVA with Sidak’s multiple comparisons test to compare heme treated and untreated cells for each sensor. Black asterisks indicate significant differences between heme treated and untreated Mtb cells: (HS1) *p* = 0.8353; (HS1-M7A) **** *p* < 0.0001; (HS1-M7A,H102A) *** *p* = 0.0001. Gray asterisks denote significant differences between HS1 and the indicated heme sensor in cells cultured with matched amounts of heme: **** *p* < 0.0001; ** *p* = 0.0015. In panel **b**, the statistical significance was assessed by two-way ANOVA with Bonferroni post hoc test. Black asterisks denote significant differences relative to day 3 for each sensor or growth condition: *** *p* = 0.0002; “ns” denotes non-significant differences, with *p* = 0.6681 and > 0.9999. Gray asterisks denote significant differences relative to HS1 for a given day: **** *p* < 0.0001. In panel **c**, the statistical significance was assessed by two-way ANOVA with Bonferroni post hoc test. Black asterisks denote significant differences relative to day 3 for each growth condition: *** *p* = 0.0004; ** *p* = 0.0029; “ns” denotes non-significant differences and *p* > 0.9999 and *p* = .0881 for day 3 vs. day 18 in untreated and ALA treated samples, respectively. Gray asterisks denote significant differences relative to WT for each time point: * *p* = 0.0358; ** *p* = 0.0087, **** *p* < 0.0001. In panels **g and h**, the statistical significance was assessed by using a two-tailed unpaired t-test: Unless otherwise noted, differences that are not statistically significant are unlabeled.

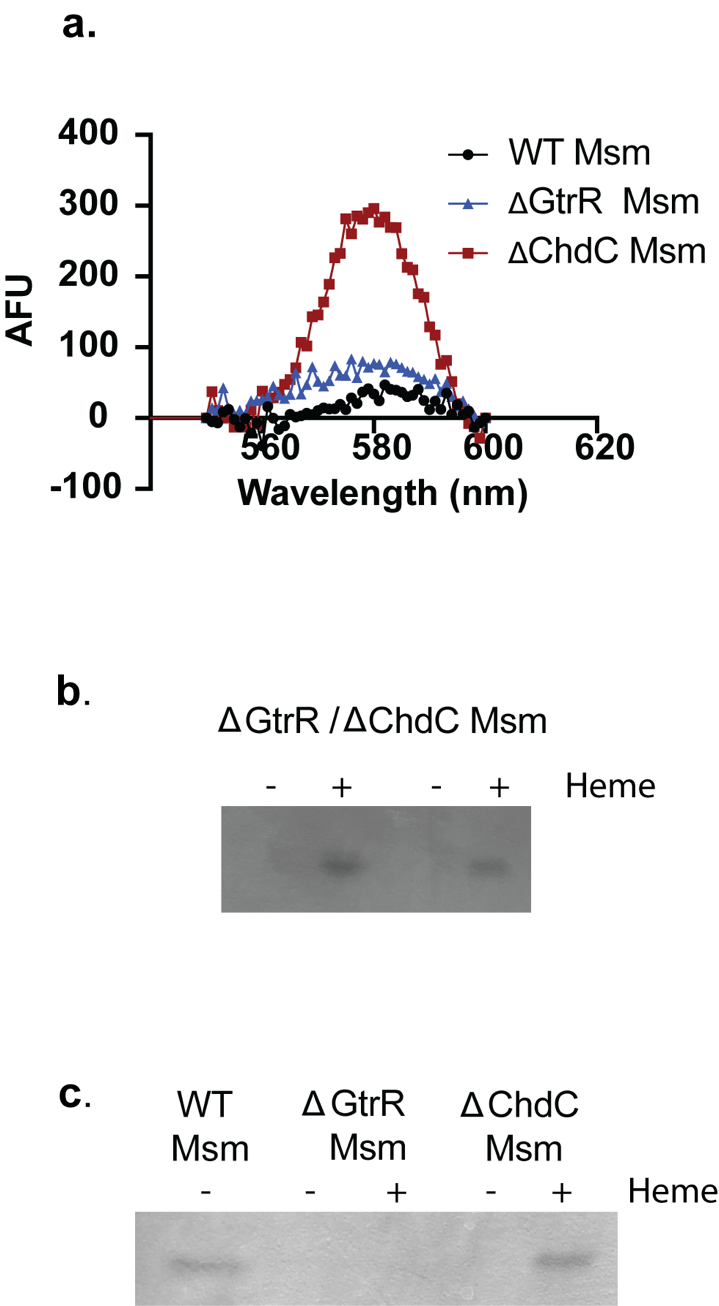

**Supplemental Figure 3. ZnMP spectra in Msm and catalase-peroxidase activity gel of ∆*gtrR*/∆*chdC* double knockout Msm**. **a**. ZnMP uptake spectra for WT, ∆*gtrR* and ∆*chdC*. Spectra are an average of 2 independent experiments. **b.** ∆*gtrR* ∆*chdC* double knockout Msm was depleted for 18 hours then treated with and without 50 µM hemin chloride for 8 hours. KATG activity is measured by in-gel catalase-peroxidase activity. **c.** Catalase-peroxidase activity gel of WT Msm without heme treatment and ∆*gtrR* Msm and ∆chdC Msm depleted (-) or with heme (+) treatment for 8 hours. Zymogram is from **Fig. S4f** and is shown to highlight differences between ∆*gtrR* Msm and ∆*chdC* Msm KatG activity after heme treatment for comparison to ∆*gtrR* ∆*chdC* double knockout. For both zymograms, dark bands indicate where KATG actively scavenged peroxide. Zymograms are shown in black and white with inverted colors for ease of viewing.

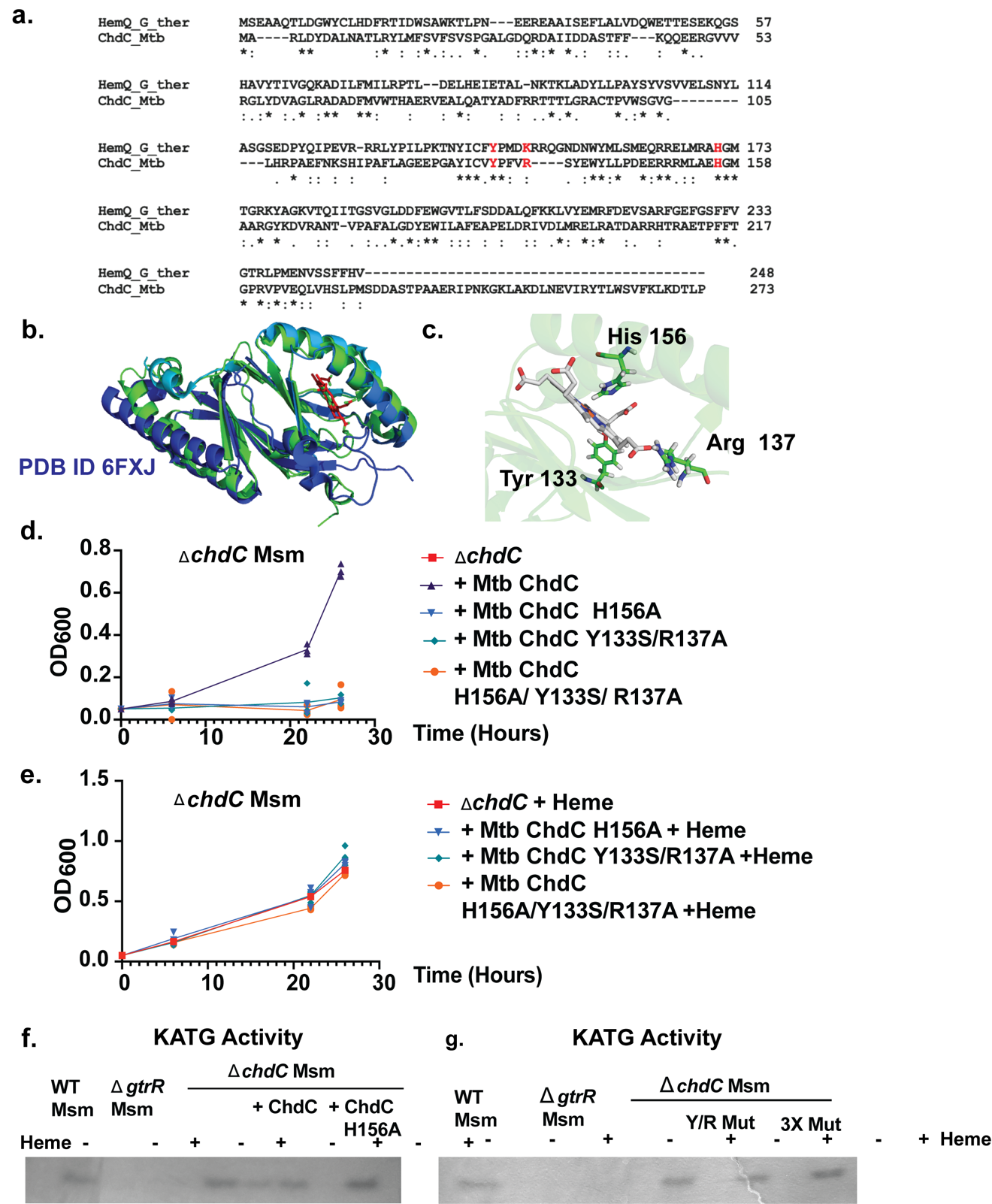

**Supplemental Figure 4. Role of ChdC catalytic activity in heme utilization. a.** The active site of Mtb ChdC was predicted from sequence alignment via Clustal Omega(16) with ChdC *Geobacillus stearothermophilus* whose active site has been characterized.(2) **b.** Homology model of Mtb ChdC in green created using the Robetta server (17, 18) with ChdC from *Listeria monocytogenes* in blue (PDB ID 6FXJ).(19) **c.** Model of active site from overlay and Robetta model in **b.** Mutated Mtb residues are shown as green sticks. Heme is from *L.* *monocytogenes* structure overlay (PDB ID 6FXJ). **d.** Growth of ∆*chdC* Msm, ∆*chdC* Msm complemented with WT Mtb ChdC or a mutant Mtb ChdC without heme supplementation or with **e** 25 µM heme supplementation. **f.** Catalase-peroxidase activity gel of WT Msm, ∆*gtrR* Msm, ∆*chdC* Msm and ∆*chdC* Msm complemented with WT or mutant Mtb ChdC grown with or without 25 µM heme. **g.** Catalase-peroxidase activity gel of WT Msm, ∆*gtrR* Msm, ∆*chdC* Msm and ∆*chdC* Msm complemented with mutant Mtb *chdC* grown with or without 25 µM heme. For (**d**) and (**e**) n=3.

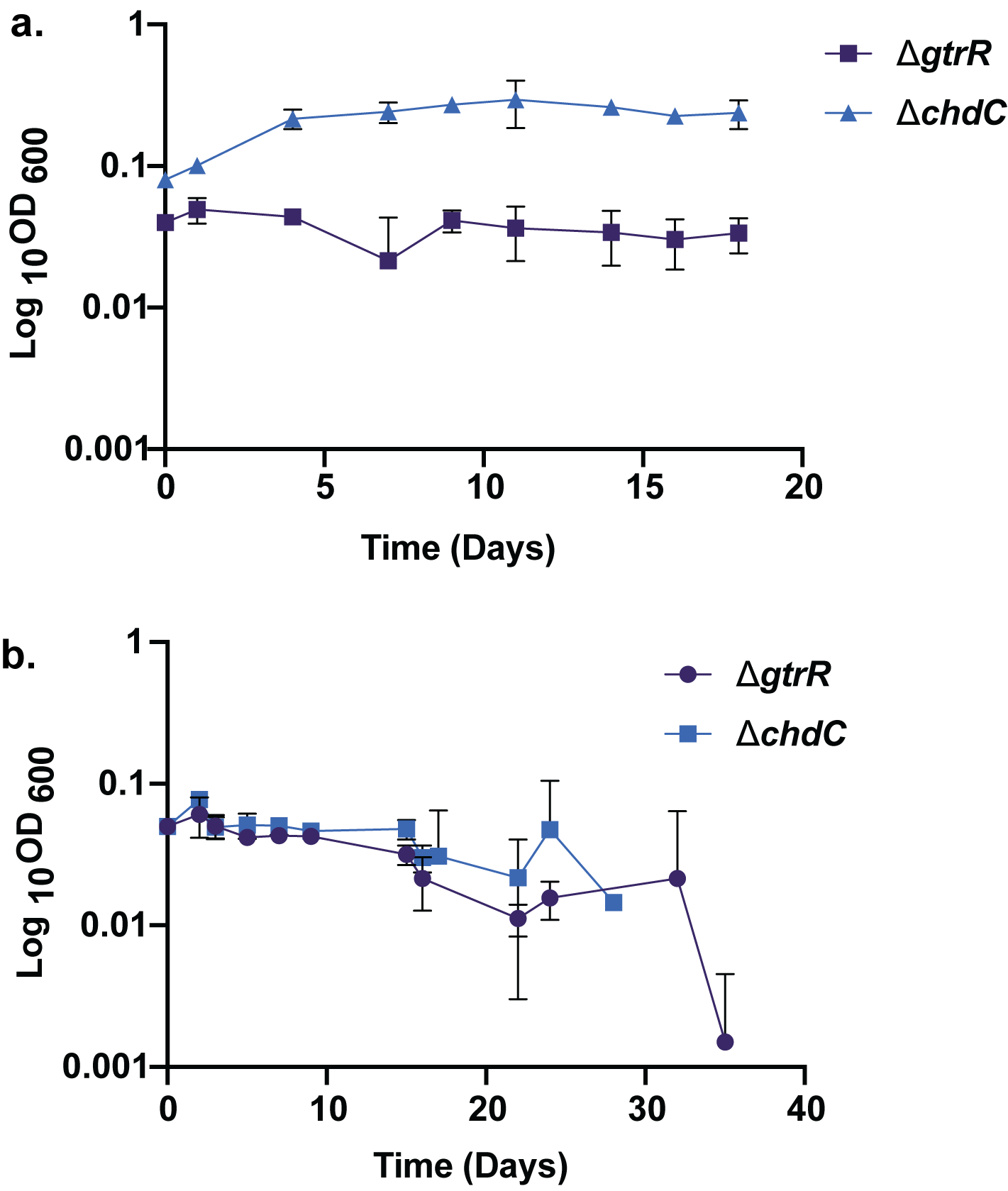

**Supplemental Figure 5. ∆*gtrR* Mtb and ∆*chdC* Mtb growth after heme depletion**. **a** After 3- day heme or ALA depletion both ∆*gtrR* Mtb and ∆*chdC* Mtb do not grow without ALA (∆*gtrR*) or heme (∆*gtrR* and ∆*chdC*) supplementation. Note In ∆*gtrR* Mtb initial OD is 0.04 and ∆chdC is 0.08. **b.** After 7- day heme or ALA depletion both ∆*gtrR* Mtb or ∆*chdC* Mtb do not grow without ALA or Heme supplementation.

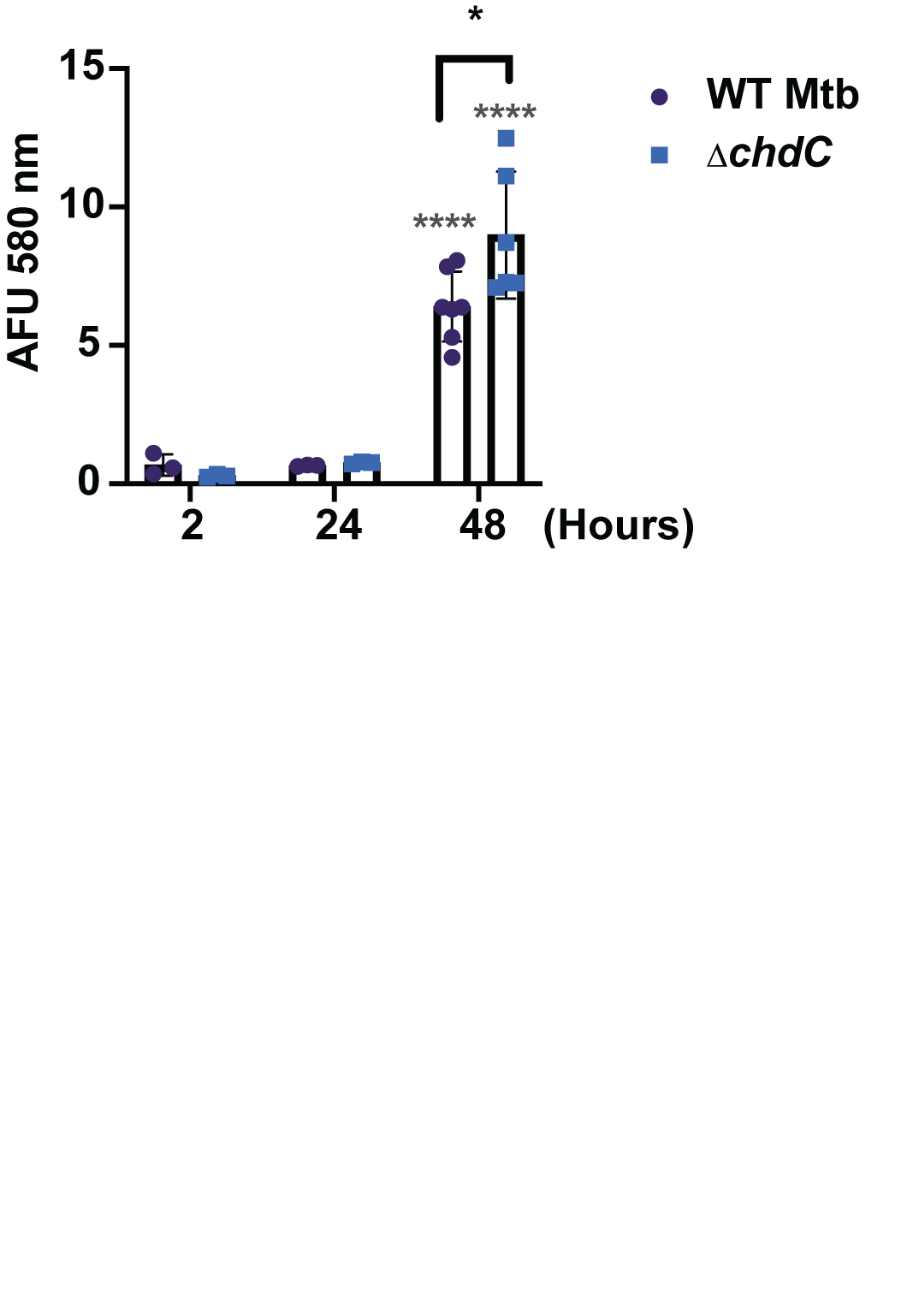

**Supplemental Figure 6. Timepoints of ZnMP uptake in *M. tuberculosis.*** ZnMP uptake measured at given timepoints adjusted to total protein of cell lysates to normalize between timepoints. P-values were calculated via a two-way ANOVA with Bonferroni post hoc test. Grey asterisks indicate P-values where ZnMP uptake is significantly increased compared to the 2-hour timepoint. *** P-value = 0.0001, **** P-value <0.0001. Black bars and asterisks indicate differences between samples at the same time point. * P-value = 0.0482. P-values between 2 hours and 24 hours for both samples were >0.9999. P=values between WT and ∆*chdC* at 2 and 24 hours were >0.9999. Error shown is S.D. for 2 and 24 hours n=3. For 48 hours WT Mtb n=7 and ∆*chdC* Mtb n=6.

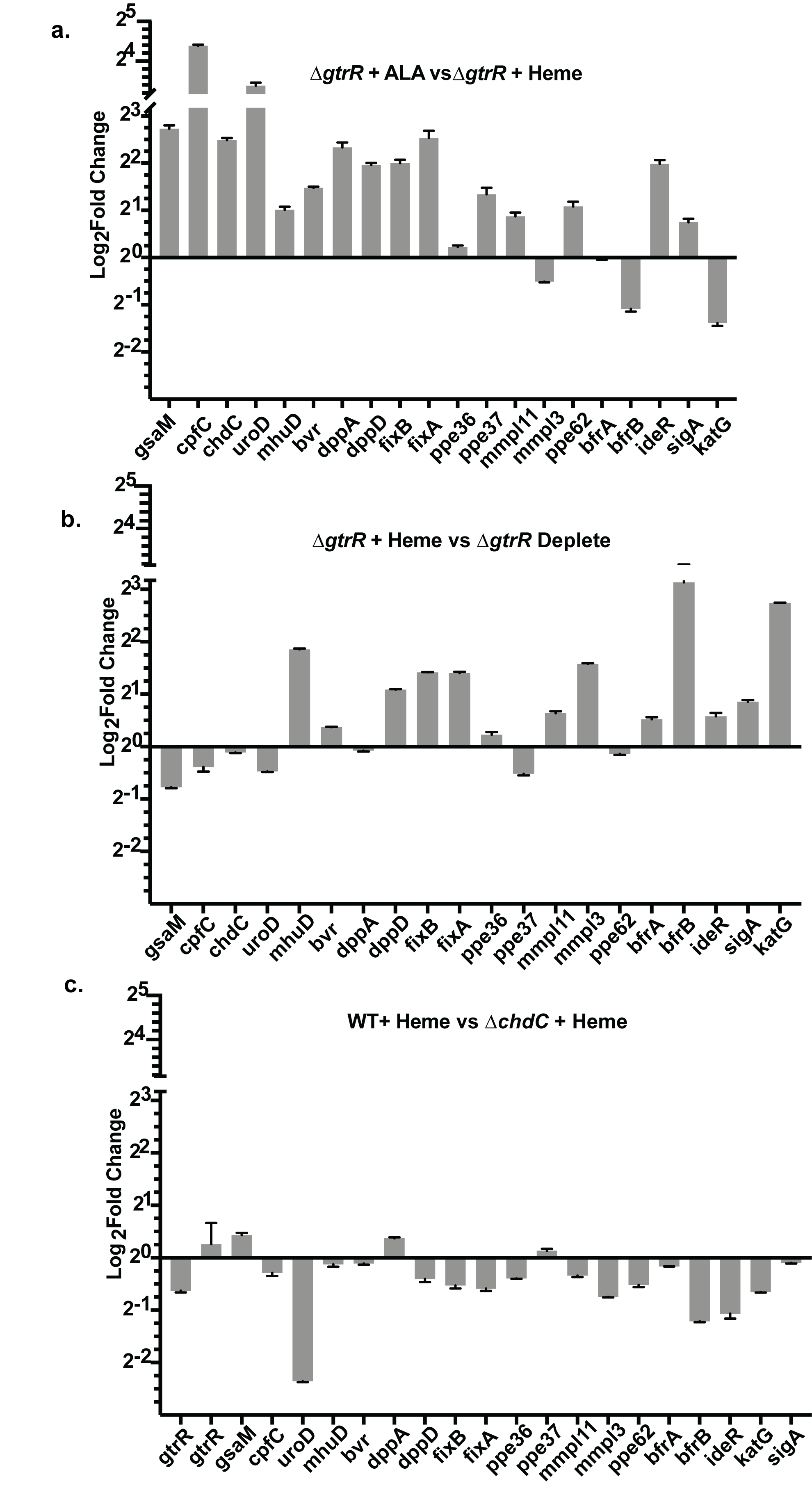

**Supplemental Figure 7. Pairwise comparisons of transcript level fold change between different growth conditions of Mtb.** All data are from two biological replicates with error bars representing +/- S.D. **a.** Log_2_ fold change between ∆*gtrR* Mtb grown in 5 µg/mL ALA versus ∆*gtrR* Mtb grown in 25 µM heme for 3 days **b.** Log_2_ fold change between ∆*gtrR* Mtb grown in 25 µM heme vs ∆*gtrR* Mtb grown in heme and then depleted for 3 days. **c.** Log_2_ fold change between WT Mtb grown in 25 Mtb grown in 25 µM heme for 3 days.

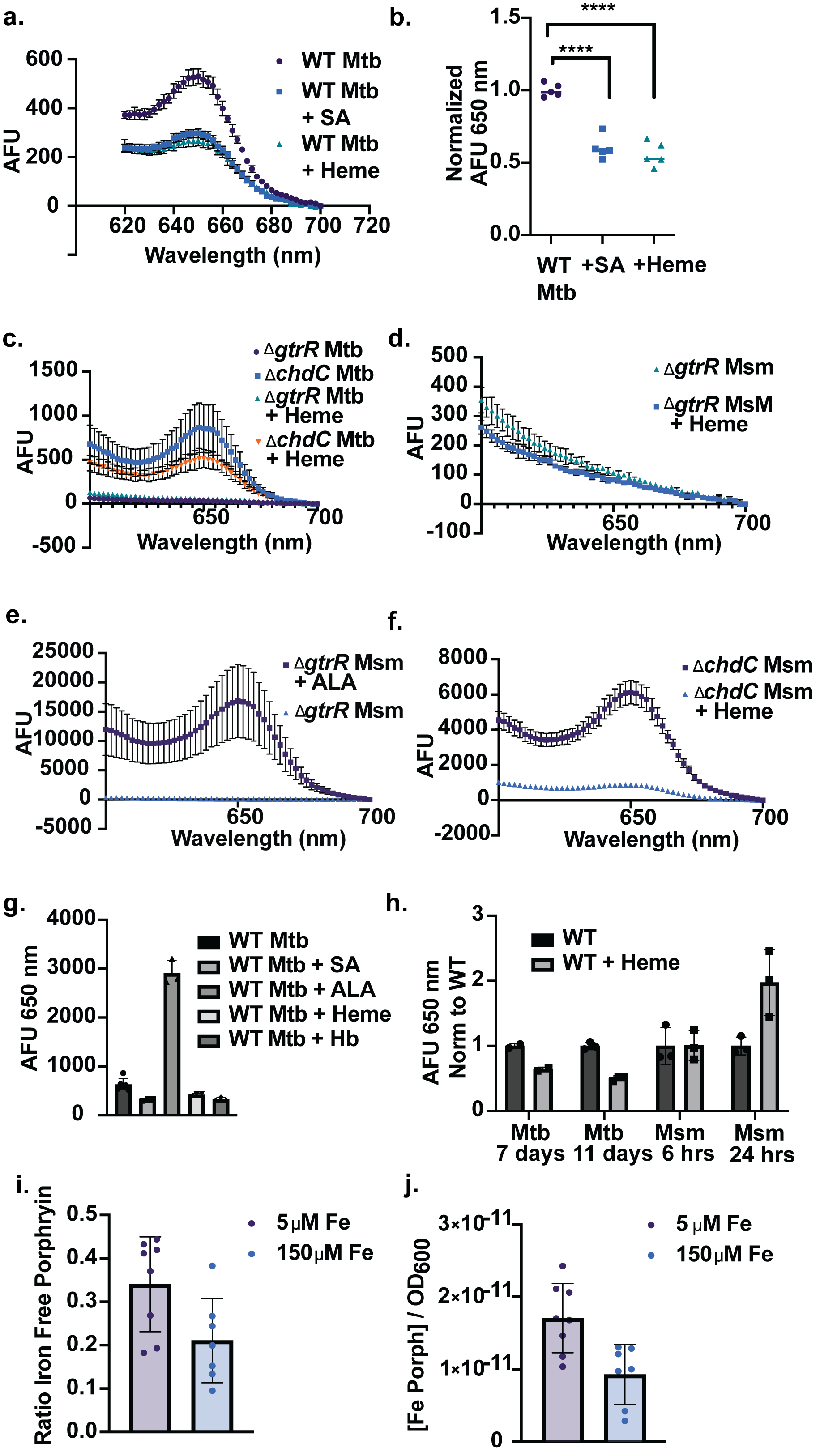

**Supplemental Figure 8. Effects of heme or iron supplementation on iron-free porphyrin accumulation in *M. smegmatis* and *M. tuberculosis*. a**. Representative iron-free porphyrin fluorescence spectra and (**b**) quantification of the emission peak at 650 nm from replicates in two independent trials are shown n=5. **c.** Iron-free porphyrin spectra of ∆*gtrR* and ∆*chdC* Mtb grown with or without 25 µM heme for one day. **c.** Iron-free porphyrin spectra of ∆*gtrR* Msm depleted for 18 hours (∆*gtrR* Msm, green circles) then after treatment of depleted ∆*gtrR* Msm with heme for 8 hours (blue circles). **e.** Iron-free porphyrin spectra of ∆*gtrR* Msm depleted for 18 hours (∆*gtrR* Msm, blue triangles)) then after treatment of depleted ∆*gtrR* Msm with ALA for 8 hours (purple circles). **f.** Iron-free porphyrin spectra of ∆*chdC* Msm depleted for 18 hours (∆*chdC* Msm, purple squares) then after treatment of depleted cells with heme for 10 hours (blue triangles). **g.** Iron-free porphyrin (FPs) levels of WT Mtb control (WT Mtb) or treated with 500 µM SA, 5 µg/mL ALA, 25 µM Heme or 6.25µM Hb for 11 days. **h.** Iron-free porphyrin fluorescence at 650 nm of WT Mtb and WT Msm treated with 25 µM Heme for given time. **i**. Ratio of iron-free porphyrin to iron-bound porphyrin (fluorescence at 650 nm in unboiled / boiled samples) of WT Mtb grown in low iron MM (5 µM Fe) and high iron MM (150 µM Fe). **j.** Concentration of iron-bound porphyrins per OD in WT Mtb grown in low iron MM (5 µM Fe) and high iron MM (150 µM Fe). Error bars shown are S.D. Spectra are average of spectra from 3 biological replicates. In (**e**) WT Mtb n=6 all other n=3. In (**f**) all are n=3 except WT Mtb 7 days where n=2. In panel **b.**, the statistical significance was assessed by a one-way ANOVA with Dunnett’s post hoc test using WT Mtb as control. In panel **b**, **** *p* < 0.0001.

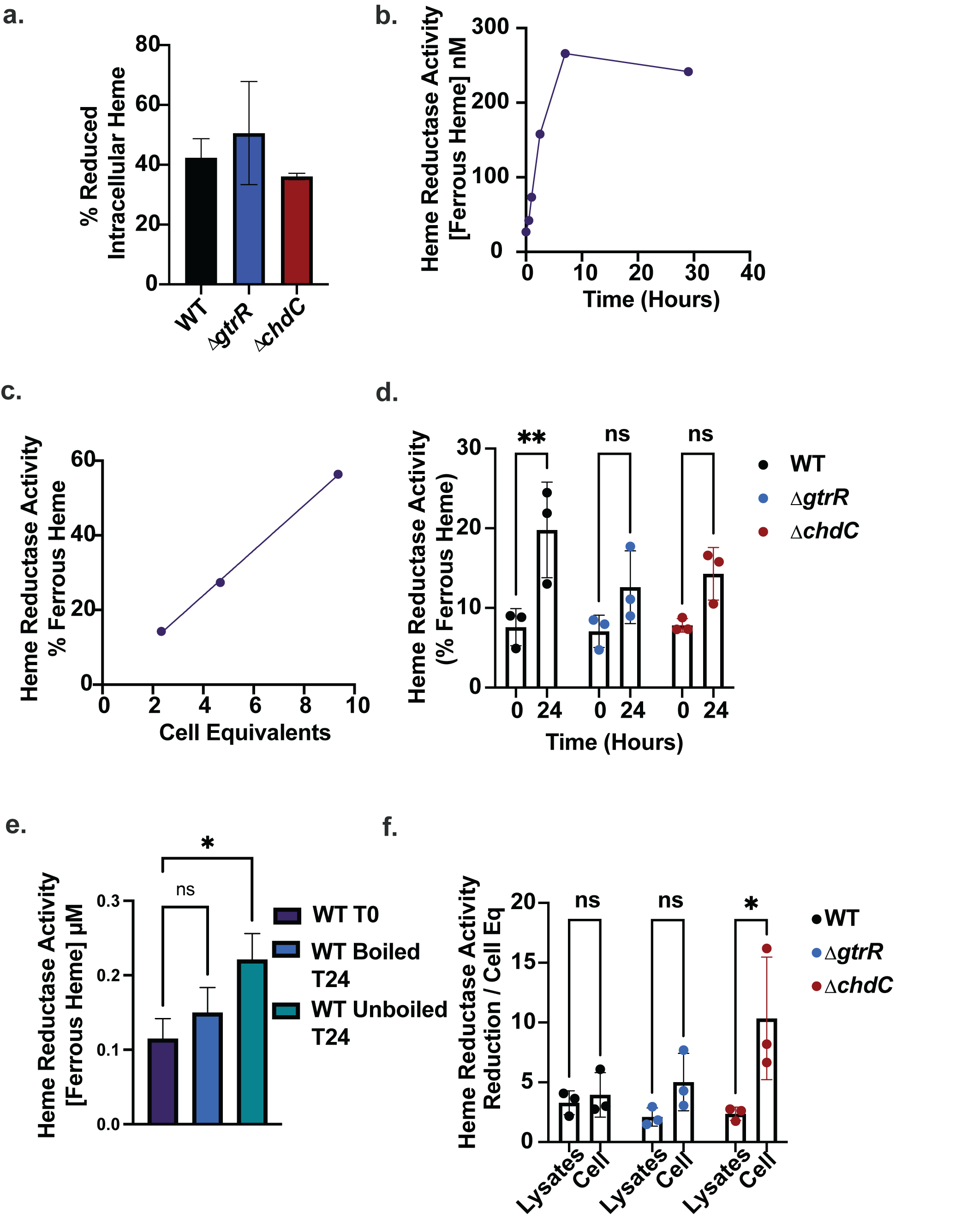

**Supplemental Figure 9. ChdC plays a role in heme reductase activity in intact cells but not disrupted cell extracts. a.** Reduced cellular heme as a percent of total cellular heme in WT, ∆*gtrR* and ∆*chdC* cells treated with 25 µM heme for 7 days. **b.** Heme reductase activity, reported as ferrous heme concentration, in WT Mtb lysates derived from a 4.7 OD_600_ pellet measured at the indicated time points. 5 µM ferric heme was added to lysates in an anaerobic chamber. **c.** Heme reductase activity, reported as % ferrous heme, in WT Mtb cells lysates derived from OD_600_ 2.3, 4.7 and 9.4 pellets after 5 µM ferric heme was added for 29 hours in an anaerobic chamber. **d.** Heme reduction in WT, ∆*gtrR* and ∆*chdC* cell lysates + 5 µM ferric heme at 0 and 24 hours in an anaerobic chamber. **e.** Heme reduction as measured by ferrous heme concentration in WT Mtb lysates immediately after 5 µM ferric heme addition (WT T0), or in boiled lysates (WT Boiled T24) and in unboiled lysates (WT Unboiled 24) 24 hours after 5 µM ferric heme addition in an anaerobic chamber. **f.** Comparison of heme reductase activities of cell lysates and intact cells in the indicated strains. Reduction of 5 µM heme in WT, ∆*gtrR* and ∆*chdC* cell lysates or intact cells after 24 hours in an anaerobic chamber was measured. To facilitate comparisons, equal cell equivalents of protein lysates and intact cells were assayed.

**Supplemental Tables**

| **Table 1: Primers used in this study** | |  |  |
| --- | --- | --- | --- |
| **Primer Name** | **Sequence** | | **Purpose** |
| HS1 Forward | GGAATTCGATATCAAGCTTTTTAAGGAGATATACATATGCACATGGTATCGGAACTGATC | | HS1 sensor plasmid construction |
| HS1 Reverse | CCCCCCCCCCCGACGTCAGGTGGCTAGCTTCATTTATACAGTTCATCCATACCCAAAGTG | | HS1 sensor plasmid construction |
| pYUB1872 Forward | ATGTATATCTCCTTAAAAAGCTTGATATCGAATTCCTGC | | HS1 sensor plasmid construction |
| pYUB1872 Reverse | CGATAGCTAGCCACCTGACGTCGGGG | | HS1 sensor plasmid construction |
| HS1 internal | AGGTGGCGGTCACTTGATTTGC | | Sequencing |
| *chdC* (Rv2676c) LL | TTTTTTTTGCATAAATTGCAGATCGTGAGCGCGTCGTC | | AES construction |
| *chdC* (Rv2676c) LR | TTTTTTTTGCATTTCTTGCGGCCCATGTTGAGCCGAATTGATGTCTCACTGAGGTCTCTACCGGGACTCACCGAGAACA | | AES construction |
| *chdC* (Rv2676c) RL | TTTTTTTTCACAGAGTGCTGCCCTTATAGAAGTGTAGCGAGTGTCTGGTCTCGTAGAGTGCCGGTGGAGCAGTTG | | AES construction |
| *chdC* (Rv2676c) RR | TTTTTTTTCACCTTGTGCACCTGGCGAAAATCGTCCT | | AES construction |
| *chdC* (Rv2676c) KoC_UP | TCGGTGGACCCGCTGTTAAG | | Deletion confirmation /sequencing |
| *chdC* (Rv2676c) KoC_DN | TCGGCCATATCGGGATTGC | | Deletion confirmation /sequencing |
| sacB out_pYUB1471 | CGGCAGGTATATGTGATGGG | | Deletion confirmation /sequencing |
| Hyg_out_pYUB1471 | AACTGCTCGCCTTCACCTTC | | Deletion confirmation /sequencing |
| *gltR* (Rv0509) LL | TTTTTTTTCCATAAATTGGCATGGTGTTCGCCGTTACCA | | AES construction |
| *gltR* (Rv0509) LR | TTTTTTTTCCATTTCTTGGGTACCAGATTCGGGATGTCTGATGTCTCACTGAGGTCTCTTACGATGCGACACCCCGAAG | | AES construction |
| *gltR* (Rv0509) RL | TTTTTTTTCCATAGATTGGGATGGTATTCACTCGGTGTCCGAGTGTCTGGTCTCGTAGGCGGATTCGACGCTGAAAGT | | AES construction |
| *gltR* (Rv0509) RR | TTTTTTTTCCATCTTTTGGGCACCGGCTCTAACGTCTCG | | AES construction |
| *gltR* (Rv0509) KoC_UP | CCTCTTTCGGGCTTTCCGTATTG | | Deletion confirmation /sequencing |
| *gltR* (Rv0509) KoC_DN | GCGACAGCTCCTCGAAGACAC | | Deletion confirmation /sequencing |
| *chdC* (MSMEG_)952) LL | TTTTTTTTCCATAAATTGGCTGGTGTTCGACGACGAGGG | | AES construction |
| *chdC* (MSMEG_)952) LR | TTTTTTTTCCATTTCTTGGCGAGCTTGGCCATAGGCCTATCG | | AES construction |
| *chdC* (MSMEG_)952) RL | TTTTTTTTCCATAGATTGGAAGCTGCCCTAGTTACCGGGTTTT | | AES construction |
| *chdC* (MSMEG_)952) RR | TTTTTTTTCCATCTTTTGGCTCATGCTGGTGGTGGTGTTCTG | | AES construction |
| *chdC* (MSMEG_)952) KoC_UP | GCGACCATCGGATTGCGTTCTG | | Deletion confirmation /sequencing |
| *chdC* (MSMEG_)952) KoC_DN | TCTACGGGTTCATGACCATGGAC | | Deletion confirmation /sequencing |
| *gltR* (MSMEG_0952) LL | TTTTTTTTCCATAAATTGGGTGCCGGGCCAGATCTTGTC | | AES construction |
| *gltR* (MSMEG_0952) LR | TTTTTTTTCCATTTCTTGGCAGCACGCTCACGGCTTCATC | | AES construction |
| *gltR* (MSMEG_0952) RL | TTTTTTTTCCATAGATTGGACTGAGTAGAGCTTGGCAAACAAC | | AES construction |
| *gltR* (MSMEG_0952) RR | TTTTTTTTCCATCTTTTGGGACCTGCCGGGTCATTTCTC | | AES construction |
| *gltR* (MSMEG_0952) KoC_UP | CGCGGTGACCAGCCAGAC | | Deletion confirmation /sequencing |
| *gltR* (MSMEG_0952) KoC_DN | GCCGAAGGTGGCCAGACTC | | Deletion confirmation /sequencing |
| Mtb *chdC* (Rv2676c) Forward | AGGAATTCGATATCAAGCTTTTTAAGGAGATATACATATGGCCCGTCTTGACTATGACGC | | Mtb WT chdC expression plasmid |
| Mtb *chdC* (Rv2676c) Reverse | ACAACGTGGCTTTCCCCCCCCCCCGACGTCAGGTGGCTAGCTCATGGCAGCGAGTGCACC | | Mtb WT chdC expression plasmid |
| Mtb *chdC* Y133S R137A Forward | CGAGGAGCCCGGCGCCTACATCTGCGTCTCTCCGTTTGTGGCGTCTTACGAGTGGTACTT | | Site directed mutagenesis |
| Mtb *chdC* Y133S R137A Reverse | ACGCCACAAACGGAGAGACGCAGATGTAGGCGCCGGGCTCCTCGCCGGCCAGAAACGC | | Site directed mutagenesis |
| Mtb *chdC* H156A Forward | CGCCGCATGCTCGCCGAAGCCGGCATGGCCGCCCGCGGATAC | | Site directed mutagenesis |
| Mtb *chdC* H156A Reverse | GTATCCGCGGGCGGCCATGCCGGCTTCGGCGAGCATGCGGCG | | Site directed mutagenesis |
| Kan pro out | TAATCGCGGCCTCGAGCAAG | | Sequencing |
| G13 Promoter seq primer | TGGTCGATACCAAGCCATTTCC | | Sequencing |

| **Table 2. Primers used for qPCR in this study.** | | | |
| --- | --- | --- | --- |
| Gene Name | Rv Number | Forward Primer | Reverse Primer |
| gtrR | Rv0509 | ACCGCCTGGCTAATGTCCTG | GCATGCCCAAGTCGCATATC |
| gsaM | Rv0524 | TTGTGCCACACCAGATTTCG | CAGCATGGCATGAAAGAACG |
| cpfC | Rv1485 | CGTTCCTGGAGAACGTTACC | ACGGCATCTTCTACATACGG |
| chdC | Rv2676c | TTTGTGCGGTCTTACGAGTG | ACGCCAGGATCCATTCGTAG |
| uroD | Rv2678c | ATTACCCTGCAGCCGATACG | TACCGGTTGAATCGCTTGTG |
| mhuD | Rv3592 | CCTCGGCTTTCAGCTGTTAC | TCAAGCACGACCTCGAATTCC |
| BVR | Rv2074 | GCGATGGTCAACACCACTAC | GGGTCGAAGGTGAAACCTAC |
| dppA | Rv3666c | CCCGTCGATGATCGAGTTTC | CGACGACACTGATGTAATCC |
| dppD | Rv3663c | GGGACTCCCGGTAAATCTTG | GTGGAGTGGTGGTGGAATC |
| fixB | Rv3029c | AAGCGCTACAGATTCGGGAG | ATGCCGTCGTCCTTTAGGTG |
| fixA | Rv3028c | TGCCGCCAAGATCTACGTC | GACACCCACTCCACCTTCTC |
| ppe36 | Rv2108 | CACGTTGCTGGAGTCGTATAG | TGCCTTCAACACTGTGGTCTG |
| ppe37 | Rv2123 | CGACCCGACCAAATTGATCC | GGCGAGAAACGTGAAGACTG |
| mmpL11 | Rv0202c | CCTGCCTATCATTCTGATGG | AGGATGAACAGGGAGTAGTC |
| mmpL3 | Rv0206c | CGGCGAATATGTGGCAAGAG | AGGCCGTGGATTGAATCCAG |
| ppe62 | Rv3533c | GAAACGGCGGCGCAATAAAC | GACCAGCCGGTATTTCTGAC |
| bfrA | Rv1876 | ACGACGTGTTGAATCGTCTC | AAGCTCCTCTCCTAGCTTGT |
| katG | Rv1908c | GACAAGGCGAACCTGCTTAC | TCCCAGGTGATACCCATGTC |
| ideR | Rv2711 | GGCTGGTCAAGGTGCTCAAC | CCCTGAACGTGCTCGGTAAG |
| bfrB | Rv3841 | GACCTTCGTGTCGAAATTCC | GATCTGTTCCTGCAAGAACC |
| sigA | Rv2703 | CAGCTGATGACCGAGCTTAG | CCTGGATCAGGTCGAGAAAC |
| 16s rRNA | rrs | CACTGGGACTGAGATACGGC | CTCCACCTACCGTCAATCCG |
